# Strategy-dependent effects of working-memory limitations on human perceptual decision-making

**DOI:** 10.1101/2021.09.03.458917

**Authors:** Kyra Schapiro, Krešimir Josić, Zachary P. Kilpatrick, Joshua I. Gold

## Abstract

Deliberative decisions based on an accumulation of evidence over time depend on working memory, and working memory has limitations, but how these limitations affect deliberative decision-making is not understood. We used human psychophysics to assess the impact of working-memory limitations on the fidelity of a continuous decision variable. Participants decided the average location of multiple visual targets. This computed, continuous decision variable degraded with time and capacity in a manner that depended critically on the strategy used to form the decision variable. This dependence reflected whether the decision variable was computed either: 1) immediately upon observing the evidence, and thus stored as a single value in memory; or 2) at the time of the report, and thus stored as multiple values in memory. These results provide important constraints on how the brain computes and maintains temporally dynamic decision variables.

## Introduction

Many perceptual, memory-based, and reward-based decisions depend on an accumulation of evidence over time (Brody & Hanks, 2016; Gold & Shadlen, 2007; Ratcliff et al., 2016; Shadlen & Shohamy, 2016; Summerfield & Tsetsos, 2012). This dynamic process, which can operate on timescales ranging from tens to hundreds of milliseconds for many perceptual decisions to seconds or longer for reward-based decisions (Bernacchia et al., 2011; Gold & Stocker, 2017), depends on working memory to maintain representations of new, incoming evidence and/or the aggregated, updating decision variable. Working memory is known to be constrained by capacity and temporal limitations (Bastos et al., 2018; Cowan et al., 2008; Funahashi et al., 1989; Oberauer et al., 2016; Panichello et al., 2019; Ploner et al., 1998; Schneegans & Bays, 2018; White et al., 1994), which implies such limitations may also constrain decision performance when the decision requires information to be maintained in working memory. Several previous studies failed to identify such constraints on working-memory-dependent decisions but used tasks involving binary choices, which may have a low sensitivity to known working-memory limitations (Liu et al., 2015; Waskom & Kiani, 2018). It remains unclear how working-memory limitations affect decisions that require interpreting and storing continuously valued quantities, representations of which are known to degrade in working-memory (Ploner et al., 1998; Schneegans & Bays, 2018; Wei et al., 2012; White et al., 1994).

To assess such effects, we examined the relationship between decision-making and working memory in the context of visuo-spatial tasks about continuous variables (visual target locations) that are sensitive to capacity and temporal limitations of working memory (Bastos et al., 2018; Funahashi et al., 1989; Panichello et al., 2019; Ploner et al., 1998; Schneegans & Bays, 2018; White et al., 1994). Specifically, we required human participants to indicate a spatial location that was informed by one or more briefly presented visual stimuli (“disks”; Fig. 1) after a variable delay. We compared the effects of variable set size and delay when the remembered location corresponded to either: 1) the perceived location (angle) of a specific disk, identified at the time of interrogation (comparable to prior studies (Ploner et al., 1998; Schneegans & Bays, 2018; Wei et al., 2012; White et al., 1994)); or 2) the computed mean angle of a set of multiple disks, which has not been examined in detail. Additionally, we examined the effects of working-memory limitations on computed locations under two conditions that are representative of certain decision-making tasks. The first was a “simultaneous” condition in which all disks (and thus all information) were presented at once. The second was a “sequential” condition in which one disk was presented later than the others. This condition required participants to adjust to a within-trial change of available decision-relevant information, typifying decisions that require evidence accumulation over time.

**Figure 1:**
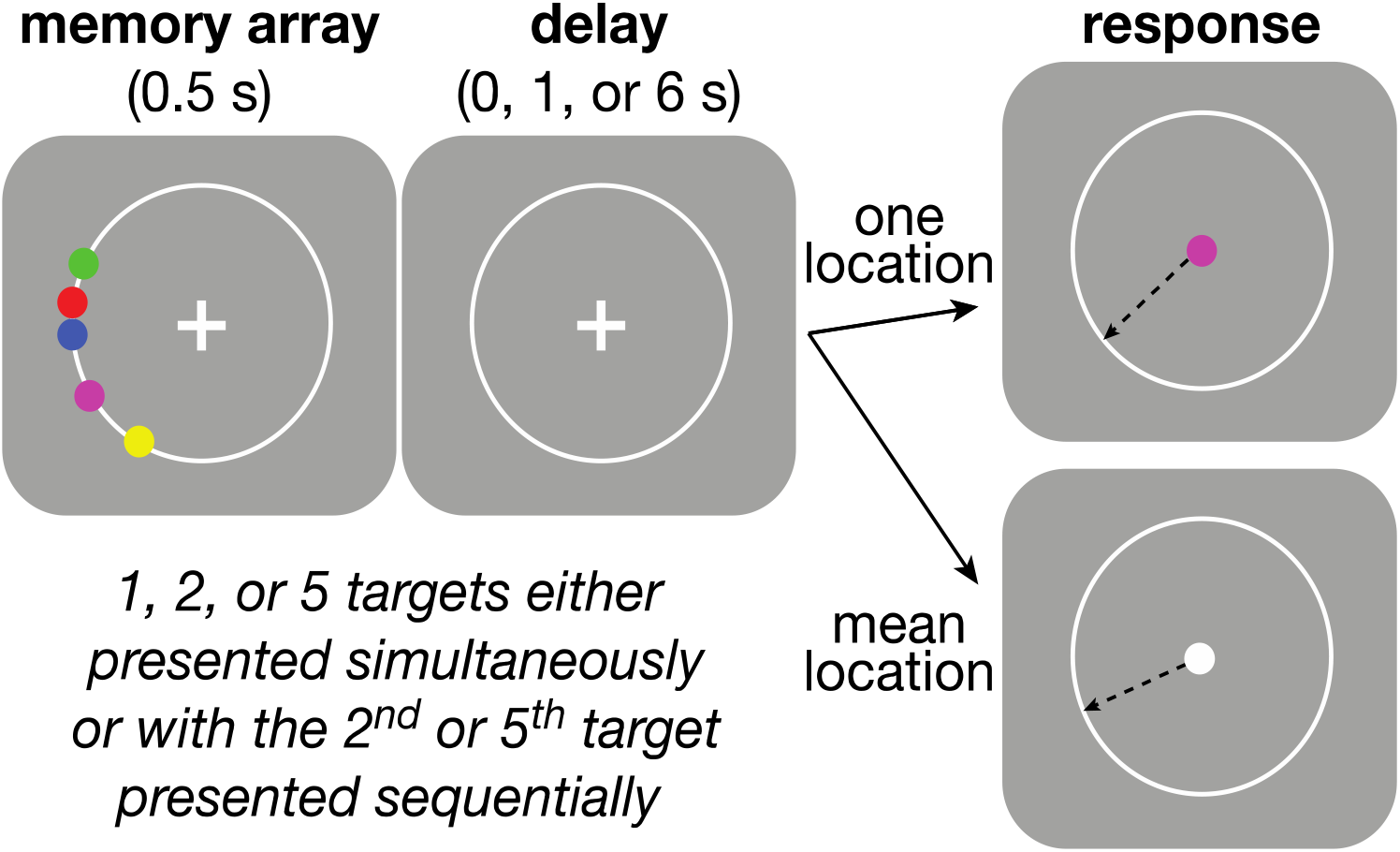
Behavioral task. Participants were asked to maintain visual fixation on the center cross while an array of colored disks was presented for 0.5 s, followed by a variable delay and finally the presentation of a visual cue that had a color that was either: 1) the same as one of the disks, indicating that the participant should use the mouse to mark the remembered location of that disk (“perceptual” trial) or 2) white, indicating that the participant should mark the mean angle of the array (“computed” trial). Perceptual and Computed trials were separated by blocks. Participants knew in advance which block they were performing, but not which disk would be probed on any given trial, during Perceptual blocks. The number of disks and length of the delay period were varied randomly within each block. Blocks were also defined by the temporal presentation of the disks. In “simultaneous” blocks all disks were presented at once, whereas in “sequential” blocks, the final disk (most counterclockwise) was presented midway through the variable delay.

For spatial working-memory tasks, the precision of working memory for perceived spatial locations is often well described by diffusion dynamics (Compte, 2000; Kilpatrick, 2018; Kilpatrick et al., 2013; Laing & Chow, 2001) that are commonly implemented in “bump-attractor” models of working memory (Compte, 2000; Constantinidis et al., 2018; Laing & Chow, 2001; Riley & Constantinidis, 2015; Wei et al., 2012; Wimmer et al., 2014) (Fig. 2a). Our analyses built on this framework by examining memory diffusion dynamics for the different task conditions and potential decision strategies. For the conditions we tested, most participants behavior was well fit by one of two strategies, each with its own constraints on decision performance based on different working-memory demands. The first strategy was to compute the decision variable (mean disk angle) immediately upon observing the evidence (individual disk angles), and then store that value in working memory in a manner that, like for the memory of a single perceived angle, could be modeled as a single particle with a particular diffusion constant (Average-then-Diffuse model; AtD; Fig. 2b parallel purple and solid black lines). The second strategy was to maintain the representations of all disk locations in working memory, modeled as separate diffusing particles, and then to combine them into a decision variable only at the time of the decision (Diffuse-then-Average model; DtA). Such strategy use results in a diffusion constant for the average that is inversely related to the number of points (Fig. 2b; magenta and dashed black lines). These two strategies had slightly different predictions and formulations when samples were presented sequentially (Fig. 2c,d). Our results show that like perceived angles, memory for computed mean angles degraded with increased set size (of relevant information) and delay between presentation and report. However, the degree of degradation depended strongly on the strategy used to compute the decision variables, implying that multiple, strategy- and task-dependent effects of working-memory should be considered in the construction of future neural and computational models of decision-making.

**Figure 2:**
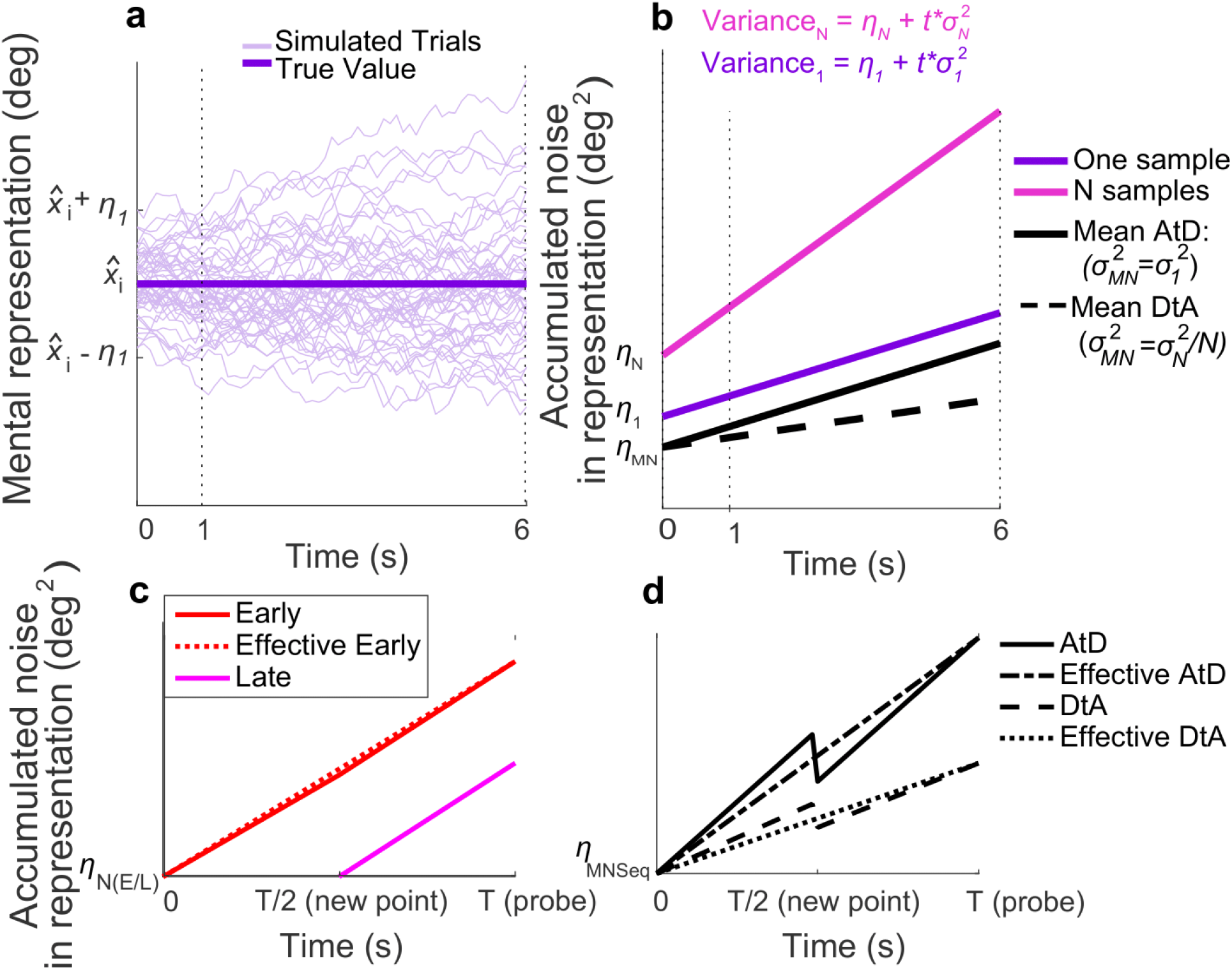
Diffusion model and predictions for different strategies. **a**) 50 simulated trials of the modeled representation of a single memorandum, 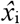, experiencing Brownian diffusion. At time *t*, the report, *r_t,1_*, is the location of the particle (motor noise is included in *η_1_*). Note the increase in variance over time, which corresponds to decreased memory precision. **b**) Linear accumulation of noise (variance) for single or multiple perceived items (colors as indicated) or computed mean values using two different strategies. In each case, the memory representation starts with some initial, additive error, *η_N_*, and diffuses over time with a diffusion constant *σ_N_^2^*, where N indicates set size. For the Average-then-Diffuse (AtD) model, the average over the presented stimuli is calculated immediately, and this single value is stored. Thus, the diffusion constant is identical between a Perceived and a Computed item (parallel purple and black lines), though the encoding may be different (i.e., the initial errors, *η_1_* and *η_MN_*, may not be equal). For the Diffuse-then-Average (DtA) model, all items are stored until the probe time. Thus, the diffusion constant of the Computed item is 1/*N*^th^ the diffusion constant of the multiple Perceived items held in memory. **c**) Accumulation of noise for Perceived items presented sequentially. Note that when the new point is added at time *T*/2, the diffusion constant for previously presented items (Early) changes slightly because of the increased load. The “effective Early” trace shows the net gain in variance over time that would be expected when sampling the error only at time T, as was done in this study. **d**) Accumulation of noise for Computed items under sequential presentation conditions for both models. At time=T/2, the final point is averaged, causing a change in the diffusion coefficients. The effective lines represent the measured change in variance over time one would measure when recording only at time T, as we did. In these examples *N*=5, *A*=0.5.

## Results

We measured the ability of human participants to remember spatial angles as a function of set size (1, 2, or 5 items), delay duration (0, 1, or 6 s), and task context (Perceived or Computed blocks). We measured error between reported and probed angles as a proxy for working memory-representations and inferred rates of memory degradation (diffusion constants) from the increase in variance of these errors over time. Below we first describe results from Simultaneous conditions, in which all items were presented simultaneously at the beginning of each trial, and show how capacity and temporal constraints on working memory relate to the accuracy of computed decision variables. We then describe our findings from Sequential conditions, in which one item was presented after the others in each trial, and show how capacity and temporal constraints affect the process of evidence integration over time.

### Simultaneous condition behavior

The difference in reports of Perceived spatial angles and the true probed location (i.e., the response error) tended to be relatively unbiased in that the mean error across participants was not significantly different from zero (Fig. 3a, full distributions in Fig. S1, individual participant mean errors in Fig. S3–4). However, the variance of these errors increased roughly linearly over time (Fig. 3c), consistent with predictions of particle diffusion models (Compte, 2000; Kilpatrick, 2018; Kilpatrick et al., 2013; Laing & Chow, 2001). This error variance also depended systematically on set size (Fig. 3c). However the change in error variance over time (slope of variance increase) did not depend on set size (ANOVA, significant effect of set size, F(2,32)=83.87, p=1.88e-13, and delay, F(2,32)=29.55, p=5.37e-08, but no significant interaction between set size and delay, F(4,64)=1.36, p=0.256).

**Figure 3:**
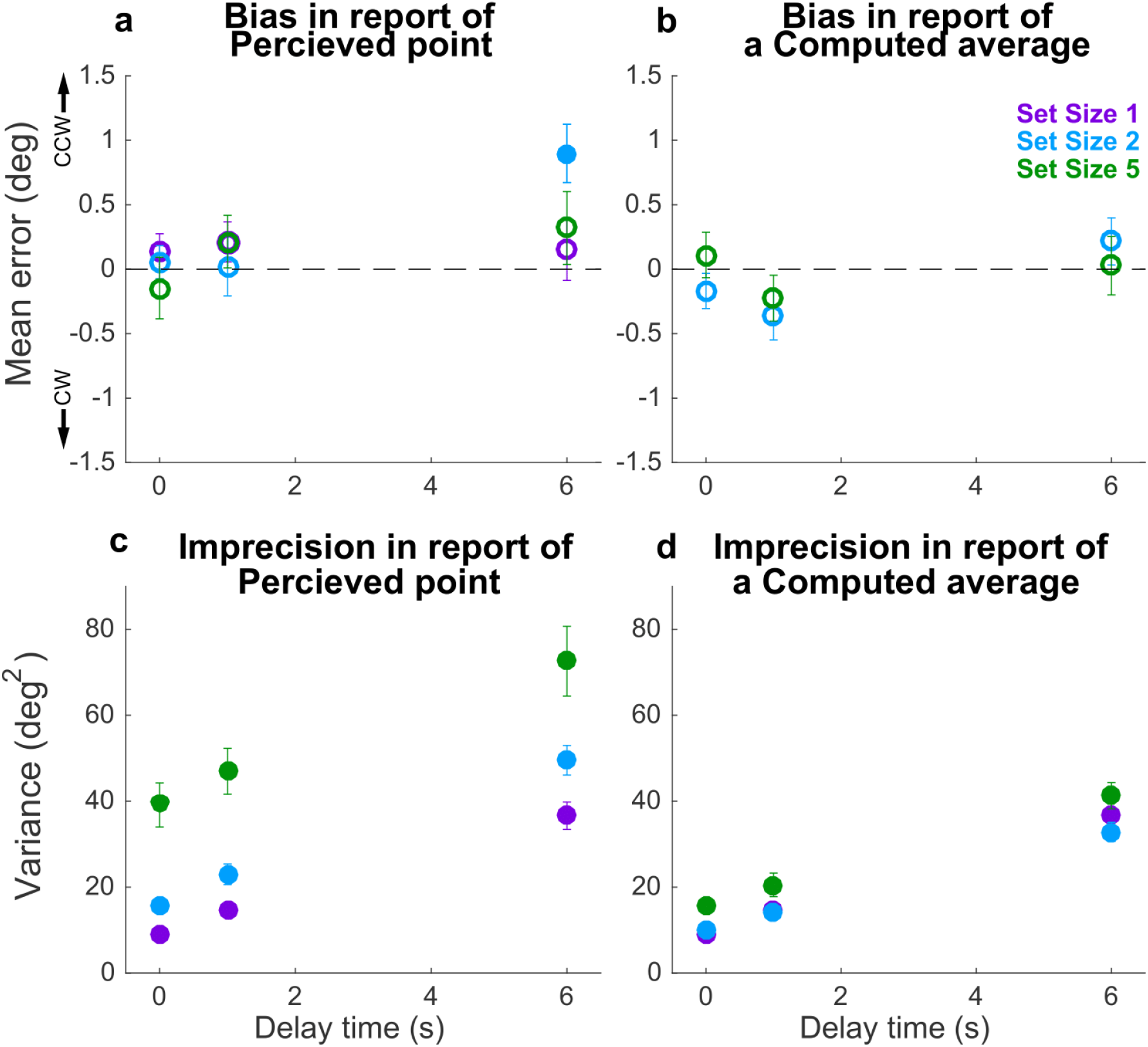
Behavioral summary for the Simultaneous condition. A) Mean Perceptual error for different set sizes (colors, as indicated) and delay time (abscissa). Filled points indicate two-tailed *t*-test for *H_0_*: mean=0, *p*<0.05. B) Mean Computed (inferred mean) error for different set sizes (colors, as indicated) and delay time (abscissa); for all tests mean error was not significantly different from zero (open circles). C) Variance in Perceptual errors, plotted as in A. D) Variance in Computed (mean) errors, plotted as in B. In each panel, points and error bars are mean±SEM across participants.

Errors in reports of Computed (i.e., inferred mean) spatial angles relative to true mean angles showed similar trends, albeit with a much weaker dependence on the number of items. Specifically, Computed angle reports were also unbiased (mean error from the true value was not significantly different from zero; Fig. 3b, S1,S3–4) but degraded (became more variable) with a roughly linear increase in variance over time (Fig. 3d). Error variance was higher at higher set sizes (set size 5 had higher variances), but the rate of degradation in accuracy did not depend on set size (ANOVA, significant effect of set size, *F*(2,32)=13.53 *p*=5.515e-5, and delay, *F*(2,32)=130.79, *p*=4.441e-16, but not their interaction, *F*(4,64)= 0.538, *p*=0.708).

### Simultaneous condition model fits

To better understand the effects of delay and set size on working-memory representations of Perceived and Computed angles for individual participants, we fit the AtD and DtA models (see Methods for details) separately to data from each condition and participant (Table 1; the two models each had the same number of free parameters and thus were compared using the log-likelihoods of the fits). Both models include terms that quantify separately the effect of set size on non-time-dependent noise (i.e., the variance in report errors with no delay; *η*) and the diffusion constant (i.e., the rate at which the variance of the errors increases over time for a single Perceived point; *σ_1_^2^*). The *A* parameter governs the relationship between *σ_1_^2^* and the diffusion constant for multiple Perceived points (*σ_N_^2^)* (i.e., *σ_N_^2^*=*σ_1_^2^*\**N^A^*).

**Table 1.** Summary of model fits for the Simultaneous condition. Parameters are: 1) ***η_1_***, non-time-dependent noise of a single value; 2) ***η_N_***, non-time-dependent noise of *N* points; 3) ***η_MN_***, non-time-dependent noise of the mean of *N* points; 4) ***σ_1_^2^***, diffusion constant of a single point; and 5) ***A***, diffusion cost of additional points. For each parameter, the maximum likelihood estimates (mean over participants±SEM) are given for the participants best fit with a particular model.

| | *Set*<br>*size*<br>*(N)* | *Number*<br>*best-fit*<br>*participants* | $\eta_I$ | $\eta_N$ | $\eta_{MN}$ | $\sigma_I^2$ | $A$ |
| --- | --- | --- | --- | --- | --- | --- | --- |
| **AtD** | 2 | 8 | 10.79±1.45 | 16.49±2.67 | 9.39±1.07 | 4.85±0.44 | 0.0892±0.24 |
|  | 5 | 14 | 9.80±1.18 | 36.88±5.33 | 14.79±1.50 | 4.34±0.47 | 0.0051±0.07 |
| **DtA** | 2 | 9 | 8.22±1.35 | 14.16±3.13 | 10.45±2.09 | 3.67±0.58 | 0.61±0.22 |
|  | 5 | 3 | 7.63±0.30 | 45.49±17.03 | 21.10±7.59 | 5.14±0.28 | 0.49±0.14 |

For the participants best fit by the AtD model, the mean, best-fitting values of *A* were close to zero, which reflects the lack of interaction between set size and delay seen in the Perceptual ANOVA in Fig. 3c (because in this model, *σ_N_^2^*=*σ_1_^2^* when *A=0)*. Conversely, for the participants best fit by the DtA model, the mean, best-fitting values of *A* were slightly higher, which reflects the lack of interaction in the Computed ANOVA in Fig. 3d (because in this model, the diffusion of a Computed point scales with *σ_N_^2^*, specifically *σ_MN_^2^*=*σ_N_^2^*/N; thus, *σ_MN_^2^* does not differ from *σ_1_^2^* in DtA only when *σ_N_^2^* > *σ_1_^2^*, which occurs when *A*>0). Of note, when *A*=1, the AtD and DtA models make identical predictions, namely 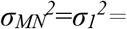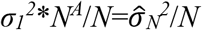. Across the population, the 95% confidence intervals for *A* (as determined by the SEM *A* values across the population) did not overlap with 1, supporting the distinguishability of the two models on average participants (although not for each individual participant; Fig. S7).

### Simultaneous condition model validation

When *A* differs from one, AtD and DtA make distinct, strong assumptions about the diffusion constant relationships between either single (AtD) or multiple (DtA) Perceived angles(s) versus a Computed average angle, as depicted in Fig. 2b. We used these assumptions to validate whether the better-fitting model and best-fit parameters for a given participant at a given set size were likely to produce the participant’s behavior. Specifically, the AtD model assumes that the diffusion constant for a single Perceived angle and for a Computed average angle are the same because both involve the memory of a single value (eq. 9). In contrast, the DtA model assumes that the diffusion constant for a Computed average angle is 1/*N*^th^ the diffusion constant for *N* points because all *N* points are held in memory prior to averaging (eq.10). We analyzed how consistent these assumptions were with the behavioral data (Fig. 4). Specifically, for each participant we fit a line to the measured error variances as a function of delay for a given set size in both Perceived and Computed blocks to estimate the change in variance over time (the empirical diffusion constant estimates: 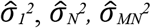, where *N*=2 or 5 for the two set sizes). We then compared the differences of these empirical estimates to the differences predicted between diffusion constants by the best fit model for a given participant.

**Figure 4:**
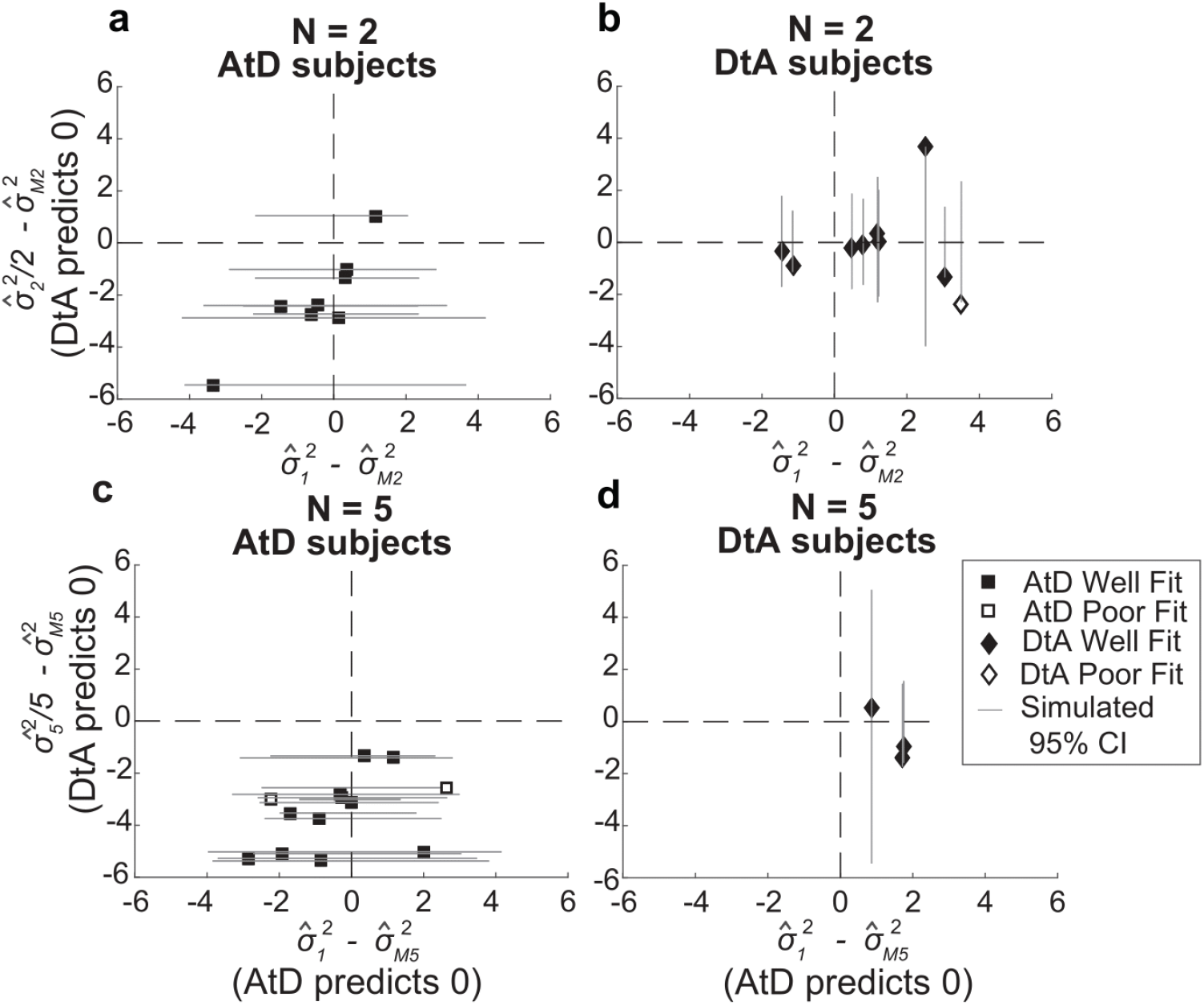
Comparisons of empirical and model-based diffusion constant relationships for the Simultaneous condition. In each panel, the abscissa shows the difference between: 1) empirical estimates of the diffusion constant for a Computed value measured by fitting a line to measured variance as a function of delay time for set size 2 (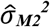, **a,b**) or 5 (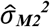, **c,d**), and 2) the empirical estimates of the diffusion constant for a single Perceived value 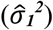. The Average-then-Diffuse (AtD) model predicts a difference of zero. The ordinate shows the difference between: 1) the empirical estimate of Computed diffusion constants 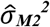 or 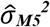, and 2) the empirical estimates of the diffusion constant for multiple Perceived values (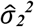 or 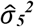) divided by the number of points. The Diffuse-then-Average (DtA) model predicts a difference of zero. Each point was obtained using data from individual participants, separated by whether they were best fit by the AtD (**a,c**) or DtA (**b,d**) model for the given set-size condition. Lines represent 95% confidence intervals computed by simulating data using the best-fit parameters for the given fit and repeating the empirical estimate comparison procedure. Closed symbols indicate participants who fell within the 95% confidence interval for their best-fit model.

In general, the participant data conformed to the model predictions of the best-fit model for that participant, despite substantial individual variability. For participants whose data were best fit by the AtD model, empirical estimates of the diffusion constant 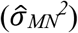 from Computed blocks tended to be similar to the empirical estimates of the diffusion constant for a single Perceptual point (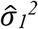; Fig. 4a,c).

Specifically, for all but two participants, the empirical diffusion constant differences fell within the 95% confidence interval of simulated distribution. Likewise, for participants whose data were best fit by the DtA model, empirical estimates of the diffusion constant 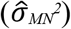 from Computed blocks tended to be similar to the empirical estimates of the diffusion constant for multiple Perceptual points divided by the set size (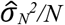; Fig. 4b,d). Specifically, for all but one participant, empirical diffusion constant differences fell within the 95% confidence interval of the simulated distribution. For some participants, the diffusion constant relationship conformed to the expectations of both models (point lying near origin in Fig. 4). These analyses thus support the idea that for most subjects, their behavior was well captured by their better-fitting model.

Summaries of the predicted report-error variances by the AtD and DtA fits for well-fit participants are shown in Fig. 5. Overall, the model predictions match participant behavior. In general, AtD participant behavior was predicted by diffusion constants that were the same for either one Perceived location or the mean Computed location based on 2 or 5 points (i.e., parallel lines in Fig. 5e,g). DtA participant behavior was well predicted by diffusion constants that were larger for multiple Perceived points compared to Single Perceived points (Fig. 5f,h). As predicted by the DtA model, the Computed point errors for DtA participants were well predicted by 1/N^th^ the diffusion constant for multiple Perceived points (Fig. 5e-h).

**Figure 5.**
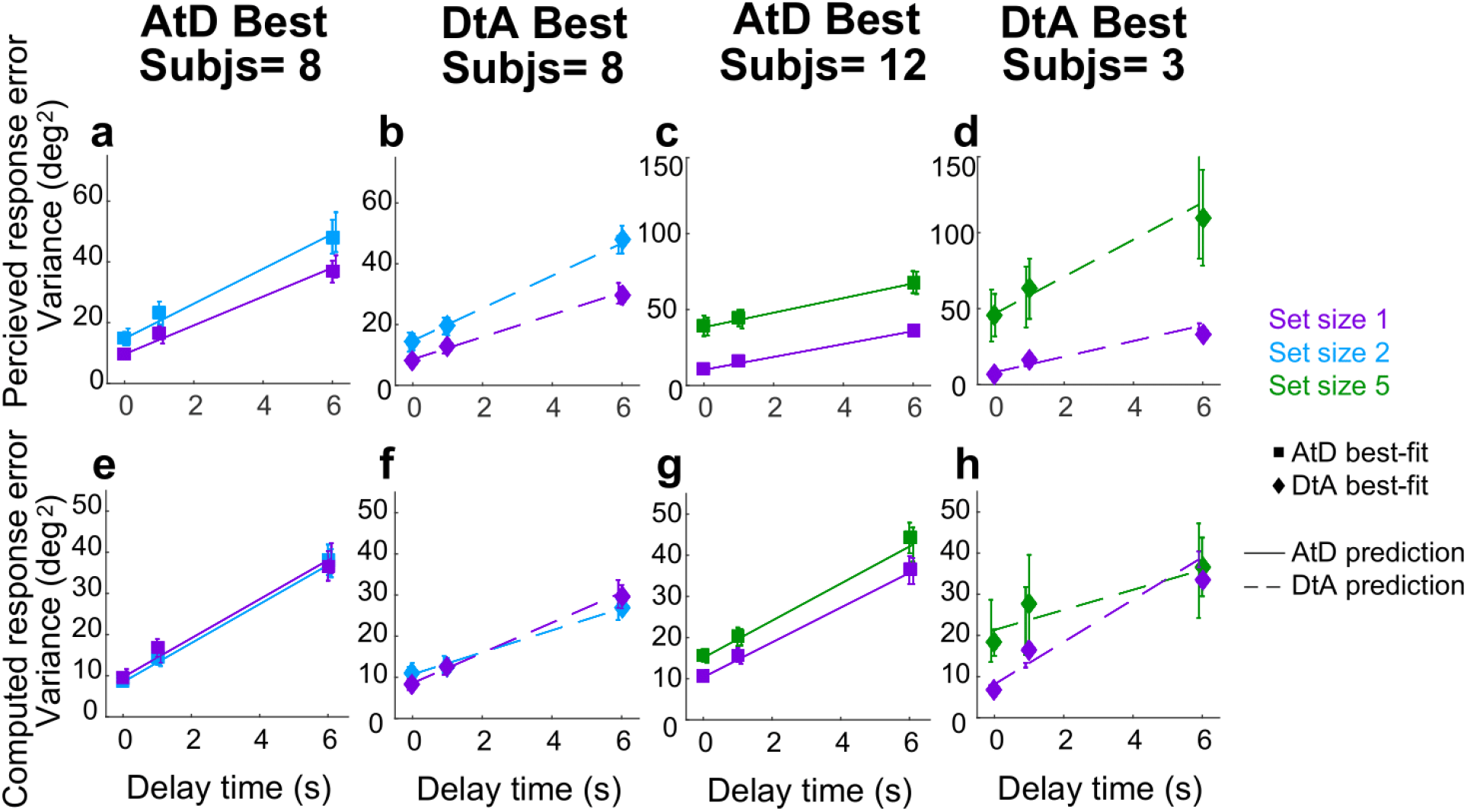
Comparison of model prediction to participant data for the Simultaneous condition. Each panel shows the empirical variance of participant errors (points and error bars are mean±SEM data across participants) and model predictions (lines, based on the mean best-fitting parameters across participants for the given model) for the participants best fit by the given model (AtD or DtA) for the given condition, as labeled above each column. A–D) Perceived blocks. E–F) Computed blocks.

### Simultaneous condition strategy comparisons

Across the population, participants had different tendencies to use the two strategies (AtD or DtA) for the two set-size conditions (Fig. 6). Specifically, equal numbers of well-fit participants were best fit by the AtD (*n*=8) and the DtA (*n*=8) model for a set size of 2, and as such neither model was significantly more likely to be a better fit (Wilcoxon signed-rank two-sided test for the median difference in the log-likelihoods of fits of the two models to data from each participant=0, *p*=0.756). In contrast, at set size 5, the well-fit participants were more likely to be better fit by the AtD (*n*=12) than the DtA (*n*=3) model (*p*=0.0027). Participants who were not poorly fit at either set size were more likely to be better fit by AtD in set size 5 compared to set size 2 (Wilcoxon signed-rank two-sided test for equal median log-likelihoods difference of fits of the two models across set sizes, *p*=0.029). Additionally, the log likelihood difference of strategy use did not correlate with the age of the participants (Pearson correlation, Fig. S8a, *p*>0.20). These findings suggest that working-memory load may affect people’s decision strategies, such that a higher load seems to correspond to an increased tendency to discard information about individual samples (disk locations) and hold only the relevant computed decision variable in memory.

**Figure 6.**
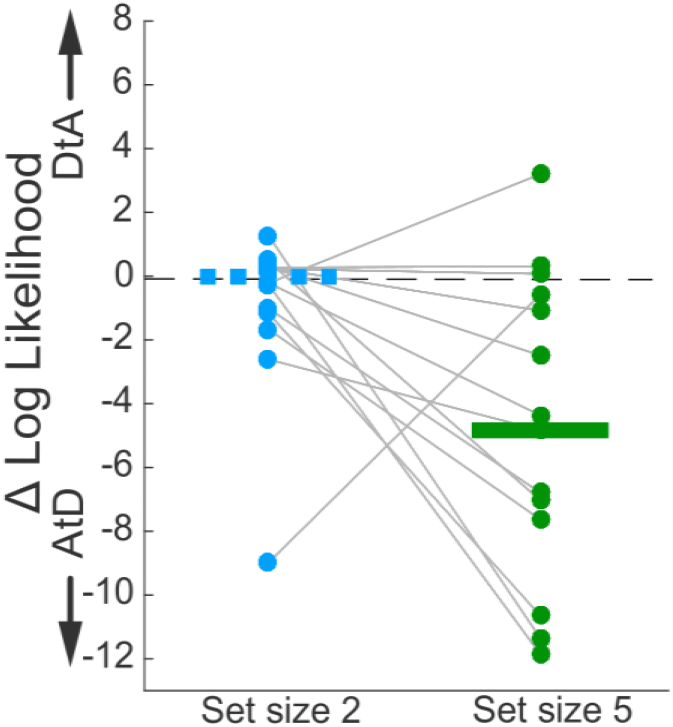
Difference in log likelihood between AtD and DtA fits for the Simultaneous condition. Negative values favor AtD. Each point represents the difference in fit log likelihoods for one participant; horizontal bars are medians (solid bar for set size 5 indicates two-sidesd Wilcoxon signed-rank test for *H_0_*: median=0, *p*=0.0027). Positive values favor DtA, whereas negative favor AtD. Grey lines connect data generated by the same participant. Only participants whose data were well matched to one of the two models (i.e., within the 95% confidence intervals depicted in Fig. 4) were included.

### Sequential condition behavior

We separately analyzed errors for Perceived reports of disks presented at the beginning (Early) or middle (Late) of a trial. Early Perceived reports tended to be relatively unbiased (mean error not significantly different from zero; Fig. 7a, full distributions in Fig. S2) but became more variable over time in a roughly linear manner (Fig. 7d), consistent with the predictions of the particle diffusion model. For higher set sizes, errors were more variable than at lower set sizes. The rate of variance increase over time did not depend on set size (ANOVA, significant effect of set size, *F*(2,32)=33.44, *p*=1.45e-08, and delay, *F*(1,16)=77.02, *p*= 1.64e-07, but not their interaction, *F*(2,32)=0.15, *p*=0.256). Late Perceived reports were likewise unbiased (mean error not significantly different from zero; Fig. 7b, full distributions in Fig. S2) and degraded in precision (i.e., increased in variance) over time (Fig.7e). However, this degradation did not depend on set size (ANOVA, significant effect of delay, *F*(1,16)=39.28, *p*=1.12e-05, but not set size, *F*(1,16)=0.90, *p*= 0.36 or their interaction, *F*(1,16)=0.0029, *p*=0.96).

**Figure 7:**
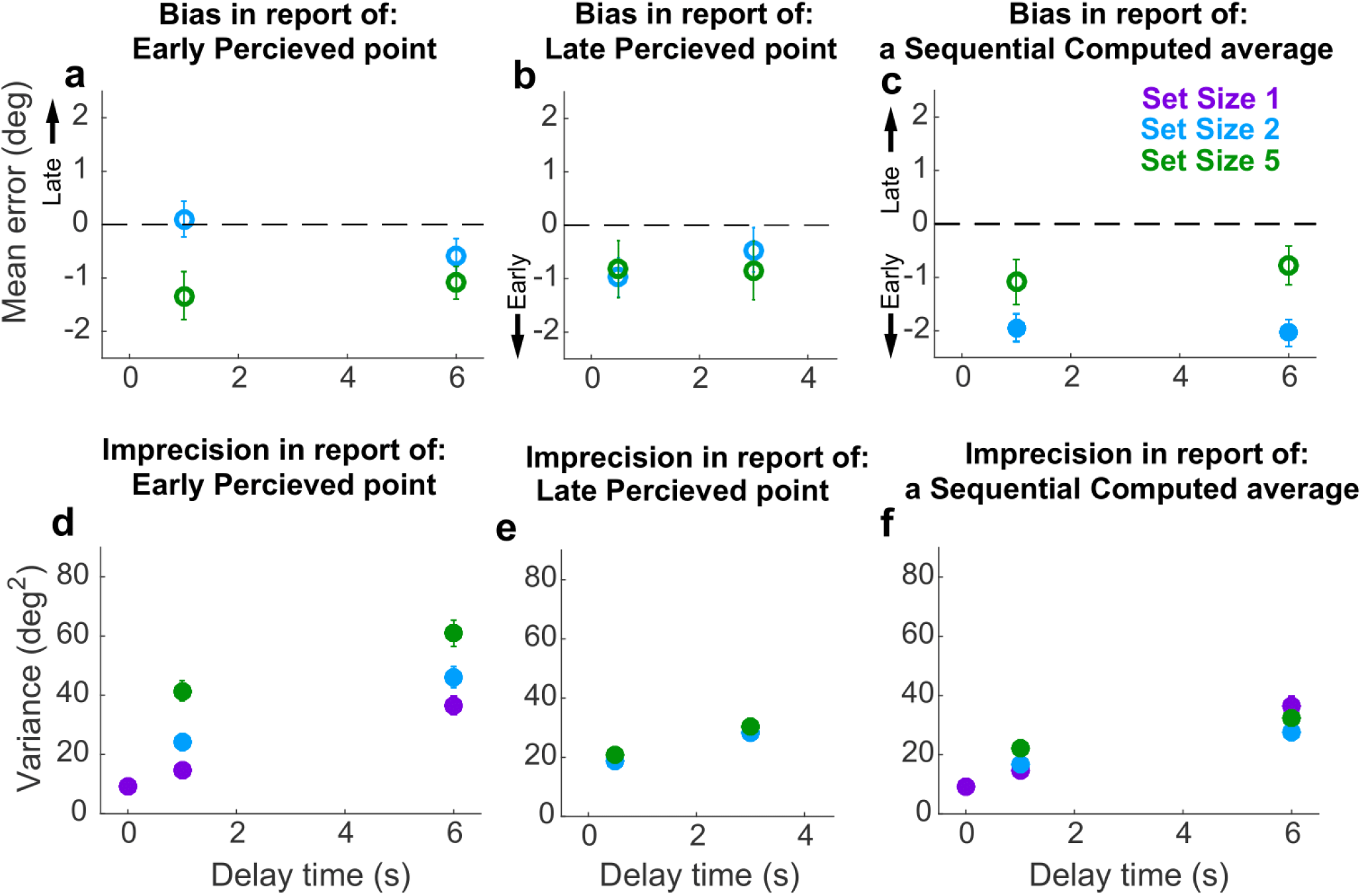
Behavioral summary for the Sequential condition. **a**) Mean error for initially presented (Early) Perceptual points for different set sizes (colors, as indicated) and delay time (abscissa). **b**) Mean error for midway presented (Late) Perceptual points for different set sizes (colors, as indicated) and delay time (abscissa). **c**) Mean Computed (inferred mean) error for different set sizes (colors, as indicated) and delay time (abscissa). Filled points in **a-c** indicate two-tailed student *t*-test for *H_0_*: mean=0, *p*<0.05. **d**) Variance in Early Perceptual errors plotted as in **a**. **e**) Variance in Late Perceptual errors, plotted as in **b**. **f**) Variance in Computed (mean) errors, plotted as in **c**. In each panel, points and error bars are mean ± SEM across participants.

Conversely, Computed (i.e., inferred mean) reports that required integrating both Early and Late points tended to be slightly biased towards the Early points for set size 2 (student two-sided *t*-test, *p*<0.001) but not set size 5 (*p*>0.5; Fig. 6c, full distributions in Fig. S2). The Computed report errors also increased in variance over time (Fig. 7f). The overall magnitude of this imprecision and its change over time depended systematically on the number of items to remember, such that more items corresponded to a slightly greater overall variance in reports at short delays, but less gain in variance over time (ANOVA, significant effect of set size, *F*(2,32)=7.73 *p*=1.8e-3, delay, *F*(1,16)=73.76, *p*=2.18e-07, and their interaction, *F*(2,32)= 6.81, *p*=3.4e-3). This interaction of delay and set size suggests the representation of the Computed value diffused in working memory with a different diffusion constant than for a single Perceived value. This interaction is consistent with predictions of both the AtD and DtA models under these conditions, though the nature of this interaction depends on the specific model, as detailed below.

### Sequential condition model fitting

To better understand the effects of delay and set size on working-memory representations of Perceived and Computed locations for individual participants under Sequential conditions, we fit the AtD and DtA models separately to data from each condition and participant (Table 2; the two models each had the same number of free parameters and thus were compared using the log-likelihoods of the fits). Recall that the *η* parameters quantify the effect of set size on non-time-dependent noise (noise when delay is zero), whereas *σ_1_^2^* is the model-based estimate of the diffusion constant for a single Perceived point.

**Table 2.** Summary of model fits for the Sequential condition. Parameters are: 1) ***η_1_***, non-time-dependent noise of a single value; 2) ***η_NE_***, non-time-dependent noise of the Early *N*–1 points; 3) ***η_NL_***, non-time-dependent noise of the Late *N^th^* points; 4) ***η_MN-seq_***, non-time-dependent noise of the mean of *N* points; 5) ***σ_1_^2^***, diffusion constant of a single point; and 6) ***A***, diffusion cost of additional points. For each parameter, the maximum likelihood estimates (mean over participants±SEM) is given for the participants best fit with a particular model.

| | *Set size (N)* | *Number best-fit participants* | $\eta_I$ | $\eta_{NE}$ | $\eta_{NL}$ | $\eta_{MN-seq}$ | $\sigma_I^2$ | $A$ |
| --- | --- | --- | --- | --- | --- | --- | --- | --- |
| **AtD** | 2 | 9 | 9.44±1.47 | 20.32±3.89 | 16.18±3.16 | 14.02±1.58 | 4.44±0.73 | -0.34±0.44 |
|  | 5 | 9 | 10.69±1.21 | 37.06±5.22 | 13.19±2.06 | 15.94±1.69 | 4.22±0.73 | -0.09±0.15 |
| **DtA** | 2 | 8 | 10.25±1.53 | 18.30±3.31 | 17.43±1.88 | 14.00±3.11 | 4.58±0.52 | -3.00±2.58 |
|  | 5 | 8 | 9.11±1.69 | 36.59±5.54 | 22.27±4.37 | 24.45±4.60 | 4.43±0.57 | -3.89±2.90 |

For participants best fit by AtD at both set sizes, the average *A* was close to zero, which is consistent with the lack of interaction between set size and delay seen in the Early and Late Perceptual ANOVA. Unlike in the Simultaneous Condition, the participants best fit by DtA had negative *A* values at both set sizes, implying that the diffusion constant for multiple Perceived items became closer to zero as the number of points increased. While counterintuitive to the concept that adding more points should increase the diffusion constant, such a negative *A* can be explained by ceiling effects: if a participant has high levels of non-time-dependent noise, they have less room to degrade while still accurately tracking the target (i.e., not having a lapse). Alternatively, the presentation of a new point may have had a stabilizing effect on the ensemble by creating directional drift towards the new point rather than random diffusion in the remaining points (Almeida et al., 2015; Wei et al., 2012), which is not inherently accounted for in any of the present models. As in the Simultaneous condition, the models make identical predictions when *A*=1. Across the population, 95% confidence intervals of *A* did not overlap with one, supporting the distinguishability of the two models; however, this difference from one was not always true for individual participants (Estimates of *A* on a participant-by-participant basis are shown in Fig. S5–6).

### Sequential condition model validation

As in the Simultaneous condition, the Sequential condition models also make predictions about the relationship between the diffusion constants of remembered Computed and Perceived values. Once again, we assessed how well participant behavior matched these assumptions, detailed in eq. 11 for AtD and eq. 12 for DtA (Fig. 8). We fit a line to the measured variances in reporting error as a function of delay for a given set size in both Perceived and Computed Sequential blocks to estimate the change in variance over time (the empirical diffusion constant estimates: 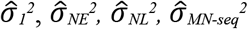, where *N*=2 or 5 for the two set sizes). We then compared the difference of these empirical estimates to the predictions of the best-fit model for each participant (Fig. 8). We used our parametric bootstrapping variant to create simulated distributions of expected deviation from the model-defined diffusion constant relationships for each subject as described previously.

**Figure 8:**
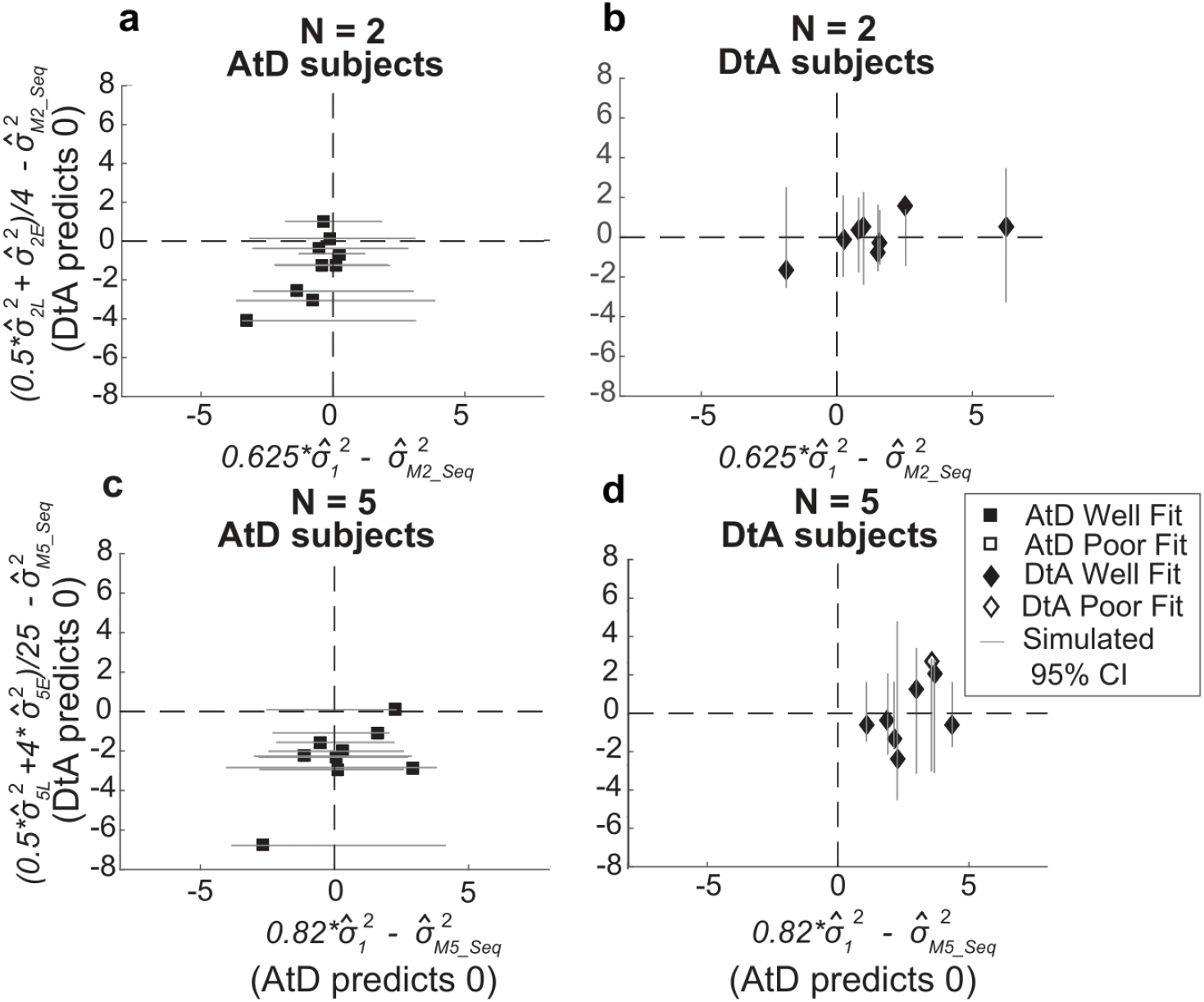
Comparisons of empirical and model-based diffusion constants. In each panel, the abscissa shows the difference between: 1) empirical estimates of the diffusion constant for a Computed value measured by fitting a line to measured variance as a function of delay time for set size 2 (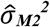, **a,b**) or 5 (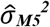, **c,d**), and 2) the empirical estimates of the diffusion constant for a single Perceived value 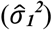 multiplied by the appropriate factor for the set size. The Average-then-Diffuse (AtD) model predicts a difference of zero. The ordinate shows the difference between: 1) the empirical estimate of Computed diffusion constants 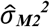 or 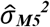, and 2) the empirical estimates of the diffusion constant of a Computed value based on the DtA hypothesis. Therefore, a value of zero indicates a match between the DtA prediction of the Computed Diffusion constant and the empirically measured estimate. Points are data from individual participants, separated by whether they were best fit by the AtD (**a,c**) or DtA (**b,d**) model for the given set-size condition. Lines are 95% confidence intervals computed by simulating data using the best-fit parameters for the given fit and repeating the empirical estimate comparison procedure. Close symbols indicate participants who fell within the 95% confidence interval for their best-fit model.

In general, the participant data conformed to the model predictions of the best-fit model for that participant, despite substantial individual variability. For participants whose data were best fit by the AtD model (n=9 for both set sizes), empirical estimates of the diffusion constant 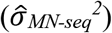 from Computed blocks tended to be similar to the expected fraction of the empirical estimates of the diffusion constant for a single point (Fig. 8a,c). Specifically, for every participant the empirical diffusion constant differences fell within the 95% confidence interval computed from simulations using the model fits. For participants whose data were best fit by the DtA model (n=8 for both set sizes), empirical estimates of the diffusion constant 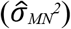 from Computed blocks tended to be similar to the expected average of empirical estimates of the diffusion constant for multiple points (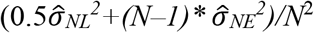; Fig. S10b,d). Specifically, for 7 participants, empirical diffusion constant differences fell within the 95% confidence interval computed from simulations using the model fits. The remaining subject was considered poorly fit and not considered in further analyses.

Summaries of the predictions of report errors variances for AtD and DtA fits are shown in Fig. 9. In general, participants best fit by AtD on average exhibited diffusion constants that were lower for Computed than Perceived values (Fig. 9i,k; lower slope of cyan/blue line versus purple line), with the difference decreasing with increased set size, as expected due to averaging process (Fig. 2d). Additionally for the participants best fit by AtD, both the Early and Late variances were on average fairly well matched by their model predictions as well (Fig. 9a,e,c,g). Conversely, participants best fit by DtA exhibited diffusion constants that were notably smaller for Computed mean locations versus single Perceived locations (Fig. 9j,l; lower slope of cyan/blue line versus purple line). The corresponding average predictions by the best fit DtA models for error variance of Early and Late Points also aligned with participant data from DtA fit participants (Fig. 9 b,f,d,h).

**Figure 9.**
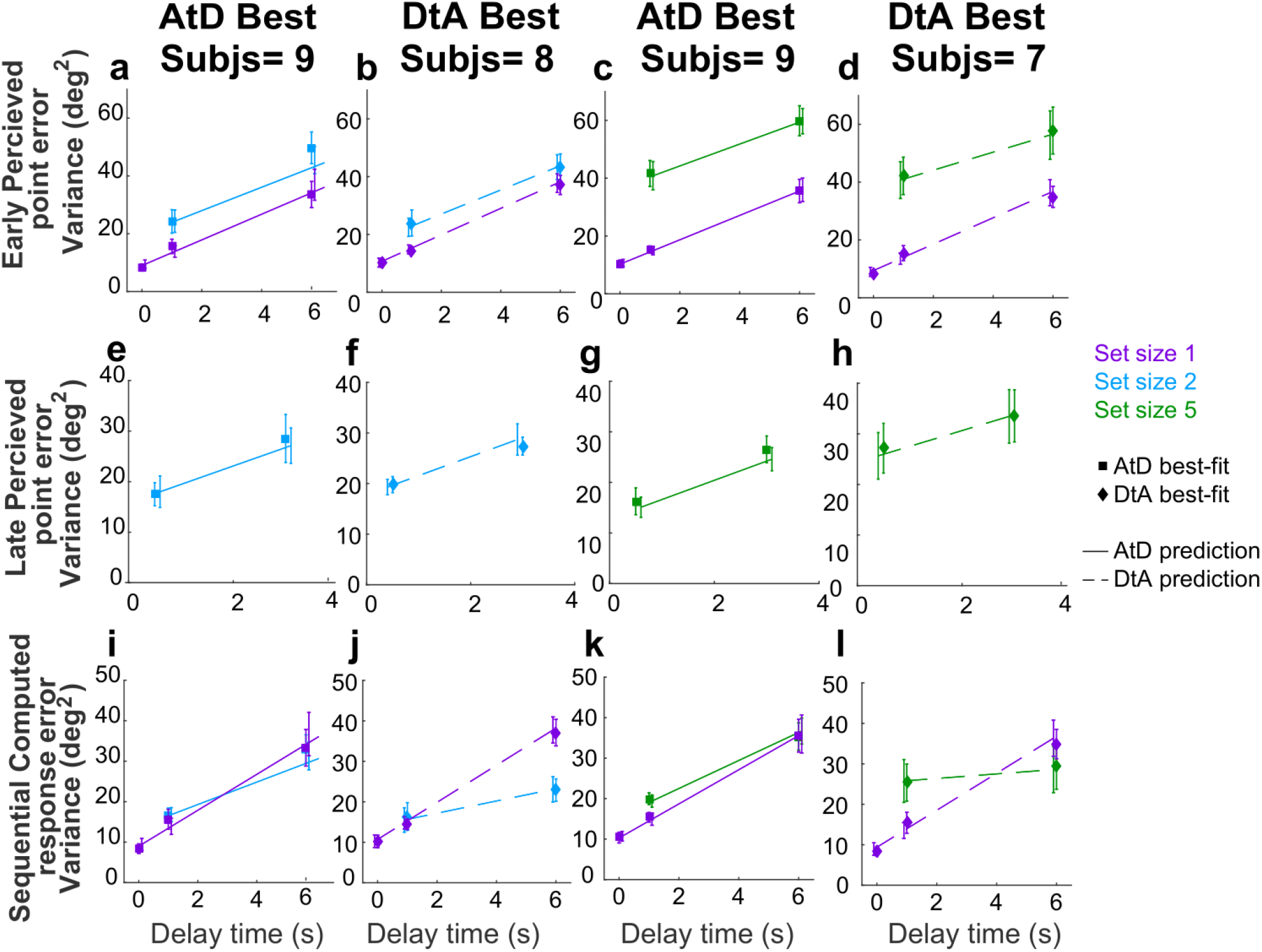
Comparison of model fits for the Sequential condition. Each panel shows the empirical variance of participant errors (points and error bars are mean±SEM data across participants) and model predictions (lines, using mean predicted variance from each participant’s best-fitting parameters for the given model) for the participants best fit by the given model (AtD or DtA) for the given condition, as labeled above each Column. Panels **a-d)** errors for Ealy points in Perceived Sequential, **e-h)** errors for Late points in Sequential Perceived blocks. **i-l)** depict errors for Sequential Computed blocks.

### Sequential condition strategy comparisons

Across the population, participants had roughly equal tendencies to use either one the two strategies (AtD or DtA) for the two set-size conditions (Fig. 10). Specifically, about an equal number of participants were best fit by the AtD (*n*=9) or DtA (*n*=8) model for a set size of 2 (Wilcoxon signed-rank two-sided test for the median difference in the log-likelihoods of fits of the two models to data from each participant=0, *p*=0.868). An approximately equal number of participants were also best fit and well fit by the AtD (*n*=9) or DtA (*n*=7) model for a set size of 5 and neither model was more likely to be the better fit across participants (*p*=0.234). Participants not poorly fit at either set size were not significantly more likely to be fit by either model across set sizes (Wilcoxon signed-rank two-sided test for identical median log-likelihoods difference of fits of the two models across set size, *p*=0.283). Additionally, the log likelihood difference of each model producing the data did not correlat e with age of participants (Fig. S8b; Pearson correlation, *p*>0.20). In general, all of the participants lost fidelity in their representation of a Computed value when it needed to consider sequentially presented information, as in many processes of evidence accumulation. However, the dynamics of this degradation differed for the two strategies, neither of which was more likely than the other for a given participant.

**Figure 10.**
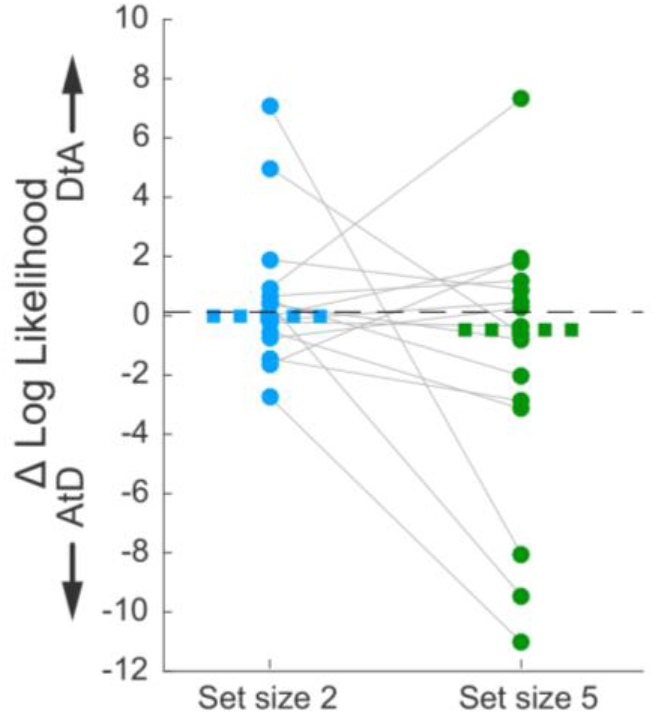
Difference in log likelihood per well-fit participant AtD and DtA fits. Negative values favor AtD. Each point represents the difference in fit log likelihoods for one participant and data from the same participant are connected across set sizes; horizontal bars are medians. Positive values favor DtA while negative values favor AtD. We failed to reject the null hypothesis (two-sided Wilcoxon signed rank test for *H_0_*: median=0, *p*>0.05) for both set sizes.

### Strategy comparisons across conditions

The use of different strategies (i.e., those captured by the AtD and DtA models) did not appear to reflect a tendency of individual participants to use a particular strategy across different conditions. Specifically, we used Fisher’s exact test of independence based on set size across temporal conditions as well as based on temporal conditions across set sizes to test whether individual participants were best fit by the same model under different task conditions. We failed to reject the null hypothesis that there is no relationship between a participant’s strategy use across set size for both Simultaneous and Sequential conditions (i.e., strategy use in set size 2 Simultaneous was not predictive of use in set size 5 Simultaneous, nor was it for Sequential conditions; *p*=0.31 and *p*=1 respectively). We also failed to reject the null hypothesis that there is no relationship between a participant’s strategy use across temporal conditions for both set sizes 2 and 5 (*p*=0.54 and *p*=1 respectively). Thus, we found that only under set size 5 were Simultaneous conditions participants more likely to use one strategy (AtD) over the other (DtA). In all other tested cases, participants were equally likely to use either strategy, and strategy use was not predictive across conditions for individual participants.

## Discussion

The goal of this study was to better understand if and how capacity and temporal limitations of working memory affect human decision-making. We used a task that required participants to report remembered spatial locations based on different numbers of objects and for different delay durations, both of which are known to systematically affect the precision of memory reports (Bastos et al., 2018; Cowan et al., 2008; Funahashi et al., 1989; Oberauer et al., 2016; Panichello et al., 2019; Ploner et al., 1998; Schneegans & Bays, 2018; White et al., 1994). We used two pairs of conditions to investigate these effects across decision-making circumstances. The first condition was Perceptual versus Computed, which allowed us to recapitulate previous findings of the effects of capacity and temporal limitations of working memory for directly observed (perceptual) quantities and then extend those findings to the kind of computed quantity that is used as a decision variable for tasks that require integration or averaging to reduce uncertainty (Brody & Hanks, 2016; Gold & Shadlen, 2007; Ratcliff et al., 2016; Shadlen & Shohamy, 2016; Summerfield & Tsetsos, 2012). The second was Simultaneous versus Sequential conditions, which extended our investigation to include the effects of working-memory limitations on decision making under relatively simple conditions (i.e., when all relevant evidence was presented at once) to the effects in a basic case of evidence accumulation over time (i.e., in which a new piece of evidence is used to update a computed quantity).

Our primary finding was that computed variables based on either simultaneously or sequentially presented information were susceptible to the same kinds of working-memory constraints as perceived variables. These working-memory limitations corresponded to a decrease in precision over time, which places critical constraints on the kinds of decision variables that are required to persist over time, such as when decisions are delayed. This result appears to contradict previous findings that found no effect of extra delays on the effectiveness of evidence accumulation for certain decisions (Liu et al., 2015; Waskom & Kiani, 2018). However, those studies used tasks with binary choices that required decision variables with less clear sensitivity to the kinds of working-memory effects we found in the context of a continuous, spatially based decision variable. Additionally, we found that increasing the number of decision-relevant points also decreased the accuracy of the continuous decision variable, although the nature of this effect was variable. More work is needed to fully characterize the conditions under which temporal and capacity limitations on the precision of working-memory representations affect decisions based on those representations.

We also found that the exact nature of interactions between working-memory limitations and decision-making depend critically on the strategy used to form the decision, and those strategies can vary substantially across individuals and tasks. For our task, we focused on two primary strategies. The first strategy, captured by the Average-then-Diffuse (AtD) model, stipulated that a participant first calculates and then stores the Computed value. Its key prediction is that a Computed value should be susceptible to the same effects of working-memory limitations as a single remembered Perceptual value in simultaneous conditions. The second strategy, captured by the Diffuse-then-Average (DtA) model, stipulated that all individual values are stored in working memory until the time of decision. Its key prediction is that the overall rate of variance increase is inversely related to the number of items. We found that participants tended to use an AtD strategy for the Simultaneous conditions with a relatively high load (five items), but otherwise were roughly equally likely to use either strategy, including for all Sequential conditions.

This finding of multiple strategy use raises several intriguing future questions. For example, we found that for the Simultaneous condition, several individuals switched from using DtA for the smaller set size to AtD for the larger set size, but we do not know if this switch was a consequence of their personal working-memory capacities. From an optimality standpoint, DtA better preserves a computed value compared to AtD for a given level of non-time-dependent noise and cost per storage item (*A)*, but only if *A* remains low (<1). It would be interesting to see if for more intermediate set sizes (i.e., 3 or 4 items) there is a reliable increase in the probability of a participant using AtD with a progression that relates to other measures of the individual’s working-memory capacity. Such future studies would more definitively support the conclusion that increased working-memory load corresponds to an increased tendency to discard information about individual samples and hold only the computed decision variable in memory. Future studies should also examine other factors that might govern which strategy is used for a given set of conditions. For example, participants in our study were instructed to report the average but given no additional details about how to do so, nor given strong incentives for choosing any particular strategy versus another. Future studies could provide more detailed instructions, incentives, and/or feedback to better understand the flexibility with which these different strategies can be employed.

Future work should also examine in more detail several other facets of working memory that were not included in our models but in principle could affect decision variables that are computed and retained over time. First, our DtA model assumed no interference between multiple items stored in memory. This assumption is undoubtedly an oversimplification, given that storage of multiple points has been both hypothesized and shown to create attraction and repulsion (Almeida et al., 2015; Kilpatrick, 2018; Wei et al., 2012). Such directional drift can create a decrease in variance over time that could affect decision variables that involve multiple quantities stored at once. Second, our DtA model also assumed that each item was stored individually. Alternatively, items could have been discarded or merged (chunked) (Kilpatrick, 2018; Wei et al., 2012), leading to different memory loads which could also affect performance. Third, we did not find strong evidence that participant behavior could not be described by the AtD or DtA model, but there was some evidence that our poorly fit subjects might have been using a strategy that started out storing multiple points as in DtA but that combined the evidence into a single variable midway through the delay in an AtD-like fashion. This kind of strategy would suggest extensive flexibility in when and how evidence is incorporated into computed decision variables, thereby placing potentially complex demands on working memory.

Both of our models were based on assumptions of a drifting memory representation. This random drift is traditionally associated with attractor models of working memory (Bays, 2014; Compte, 2000; Macoveanu et al., 2007; Wei et al., 2012) that have been used extensively to describe the underlying neural mechanisms (Funahashi et al., 1989; Shafi et al., 2007; Takeda & Funahashi, 2002; Wimmer et al., 2014). In these models, neural network activity is induced by an external stimulus and then maintained via excitatory connections of similarly tuned neurons and long-ranged inhibition. Random noise causes the center of this activity (which represents the stimulus) to drift in a manner that, dependent on the implementation, can depend on the delay duration, set size, and/or their interaction (Almeida et al., 2015; Bays, 2014; Koyluoglu et al., 2017). A recent implementation even can naturally compute a running average based on sequentially presented information (Esnaola-Acebes et al., 2021). Our results imply that such models should be extended to support the flexible use of different strategies that govern when and how incoming information is used to form such averages. It will be interesting to see if such a flexible model can account for neural activity in the dorsolateral prefrontal cortex (dlPFC), which includes neurons with persistent activity that has been associated with both spatial working memory^11,22–25^ and the formation of decisions based on an accumulation of evidence (Curtis & D’Esposito, 2003; H. R. Heekeren et al., 2006; Hauke R. Heekeren et al., 2008; Kim & Shadlen, 1999; Lin et al., 2020; Philiastides et al., 2011).

In conclusion, we found that in this spatial, continuous task, participant accuracy for both perceived and computed values was subject to working-memory limitations of both time and capacity. Additionally, we found behavior that was consistent with both the storage strategies we investigated. The fact that different participants employed different strategies for storing a computed value (such as a decision variable) and that these strategies have different consequences on overall accuracy has important implications for not only future neural network models of working memory, but also for future computational models of decision-making.

## Materials and Methods

### Human psychophysics behavioral task

We tested 17 participants (4 male, 12 female, 1 chose not to answer; age range=22–87 yrs). The task was created with PsychoPy3(Peirce et al., 2019) and distributed to participants via Pavlovia.com, which allowed participants to perform the task on their home computers after providing informed consent. These protocols were reviewed by the University of Pennsylvania Institutional Review Board (IRB) and determined to meet eligibility criteria for IRB review exemption authorized by 45 CFR 46.104, category 2.

Participants were instructed to sit one arm-length away from their computer screens during the experiment and to use the mouse to indicate choices. Each participant completed 1–2 sets of 4 blocks of trials in their own time.

The basic trial structure is illustrated in Fig. 1. Each trial began with the presentation of a central white fixation cross (1% of the screen height). The participant was instructed to maintain fixation on this cross when not actively responding. The participant began each trial by placing the mouse over the cross and clicking, to allow for self-pacing and pseudo-fixation. Initiating a trial caused a white annulus of radius 25% of the screen height to appear. A block-specific memory array appeared 250 ms later, centered at an angle chosen uniformly and at random on the anulus. The array consisted of 1, 2, or 5 colored disks sized 1.5% screen in diameter. The angular difference between any two adjacent disks was at least 6°, and between the two most distal disks was at most 60°. The disks from clockwise to counter-clockwise were always presented in the same order: green, red, blue, magenta, and yellow. When fewer than five disks were presented, the latter colors were omitted. The consistent color ordering was intended to reduce errors caused by mis-binding of location and color. The angular differences between disks in an array was randomly selected from 5 preselected sets. The memory array remained on the screen for 0.5 s, while the annulus remained on the screen throughout the delay of 0, 1, or 6 s. At the end of the delay, the fixation cross was replaced with a response cue that either matched a color of a disk in the memory array, indicating a response to the remembered location of that disk, or was white, indicating a response to the mean angle of all disks in the present trial. The response type varied by block (see below). The participant then moved the mouse and clicked on the annulus at a position at which they remembered the requested response. Feedback was then given indicating the correct location, the participant’s response, and the difference between the two.

We used four block-wise conditions: 1) Simultaneous Perceived blocks used arrays of 1, 2, or 5 disks presented simultaneously at the beginning of the trial. Participants were told in advance that they would always be asked to report the location of one of the array disks but were not informed which one until the response period. The probed disk was picked randomly on each trial. 2) Simultaneous Computed blocks used arrays of 2 or 5 disks presented simultaneously at the beginning of the trial. Participants were told in advance they would need to report the average angle of all disks shown in the present trial. 3) Sequential Perceived blocks were identical to Simultaneous Perceived blocks, except only arrays of 2 or 5 disks were used, and all but one of the disks (the counter-clockwise most) was presented at the beginning of the trial. The final disk was presented for 0.5 s ending midway through the delay of 1 or 6 s. Participants were told in advance that the final disk would be presented in the middle of the delay for these blocks. 4) Sequential Computed blocks were identical to Simultaneous Computed blocks, but with delayed presentation of the final disk as in Sequential Perceived blocks. Again, participants were told in advance that the final disk would be presented in the middle of the delay.

All participants completed one and most (12) participants completed two blocks of each type. Each block contained 50 trials at each set size and each delay time, the order of which was randomized.

### Basic analyses

Trials were excluded from analysis if the response was >30° from the correct angle. This cutoff was based on assessment of the error distributions (Fig. S1–2); using a cutoff of 25° did not noticeably change the results. On average, <10% of trials were excluded per delay condition per set size per block (see Fig. S1–2). These trials were excluded to focus analysis on trials that were directed towards the correct location and avoid lapses of attention and extreme motor errors. We investigated both the bias and variance in participant responses, as follows.

We quantified bias as the mean error between the response and the true probed angle for each participant and condition (positive/negative values imply errors that were systematically counterclockwise/clockwise respectively). A Bonferroni-corrected two-sided *t*-test was used to assess whether this mean response error was significantly different from zero across participants for each set size, delay, response type and temporal presentation. Additionally, the mean error and confidence interval for each subject were calculated for each condition (Fig. S3–6).

We quantified the variance of the error between the response and the true probed angle for each participant and condition. We chose variance as opposed to other measures of dispersion for consistency with our particle models (see below) in which variance scales linearly with delay. We examined effects of set size, delay duration, and task context on response variability using a two-way repeated measures ANOVA. On Simultaneous Perceived and Computed blocks, we used a 3 (delay duration: 0, 1, or 6 s) x 3 (set size: 1, 2, or 5 disks) within-participant design. On Sequential Perceived blocks, we used a 2 (delay duration: 1 or 6 s) x 3 (set size: 1, 2, or 5 disks) within-participants design for stimuli presented at the beginning of the trial (Early) and a 2 (delay: 0.5 or 3 s) x 2 (set size: 2 or 5 disks) design for stimuli presented halfway through the trial (Late). On Sequential Computed blocks, we used a 2 (delay duration: 1 or 6 s) x 3 (set size: 1, 2, or 5 disks) within-participants design. When the comparison included set size=1, data were always taken from the Simultaneous Perceived block.

### Model-based analyses

Our models were based on principles of working memory that are well described by bump-attractor network models (Compte, 2000; Laing & Chow, 2001; Wimmer et al., 2014). In such models, stimulus location is represented by a “bump” in activity from neurons tuned to that and similar locations. These neurons recurrently activate each other, maintaining a bump of activity even after stimulus cessation. However, because of the stochastic nature of neural activity and synaptic transmission(Faisal et al., 2008), there is variability in which neurons have the most activity at any given time (and thus are the center of the bump representing the stimulus). This variability in bump center corresponds to variability in the location representation and a degradation of the memory representation over time. The dynamics of this bump can be described as a diffusion process that obeys Brownian motion (Compte, 2000; Kilpatrick, 2018; Kilpatrick et al., 2013; Laing & Chow, 2001). We used this simplified description in our models as follows.

### Perceived values in working memory

A single point (i.e., the central spatial location of a single disk), *x_1_*, is assumed to be represented in working memory by 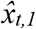, where *t* represents the time since the removal of the stimulus. We assume that 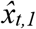 evolves like a sample from a Brownian-motion process. Specifically, when *x*_1_ is observed, it is encoded with some perceptual noise, *η^p^*. Therefore, at time zero, 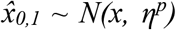. This representation accumulates noise over time with some diffusion constant, *σ_1_^2^*, further degrading the representation of 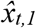 from *x_1_* such that 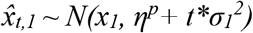. There is additional motor noise in the participant’s report, *r_t,1_*, and we denote the variance of this motor noise by *η^m^*. Mathematically, it is equivalent to add the motor noise at the beginning or the end of the diffusion of 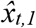 when considering the report, *r_t,1_*. In our model, we thus represent the sum of the perceptual and motor noise as a single, *non-time-dependent* noise term. Hence, we show simulated trajectories of 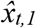 in Fig. 2a with an initial variance of *η_1_*= *η^p^+ η^m^* so that at time *t* the report, *r_t,1_* is the current angle of the trajectory. Therefore:

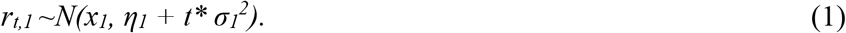

When multiple items are held in memory, they are held with less fidelity than a single point (Bays et al., 2009; Brady & Alvarez, 2015; Koyluoglu et al., 2017; Wei et al., 2012). We therefore assume that the sum of the initial perceptual noise and final motor noise, with variance denoted by *η_N_*, can depend on the number of disks, *N*. Moreover we describe, 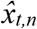, the representation of the *n*^th^ item at time *t*, by a normal distribution with a diffusion constant that is potentially higher than for a single point. We assume that this new diffusion constant σ_1_^2^, equals *σ_1_^2^*N^A^* and thus scales as a power of the total number of stimuli, *N*, held in memory (Bays et al., 2009; Bays & Husain, 2008; Wei et al., 2012), and is proportional to the diffusion constant corresponding to a single stimulus representation, *σ_1_^2^*. Therefore:

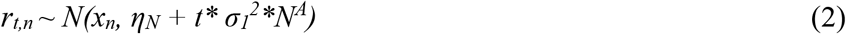

All representations in a set of size *N* share the same magnitude of non-time-dependent noise, *η_N_*, but the evolution of each representation is assumed to be independent. To examine distributions of responses across the various presented locations, we measured the error of the response *r_t,n_* relative to the true location of the target the observer was asked to report, *x_t,n_*. According to our model, the difference between the true and reported location (the error, *e_t,n_*) is

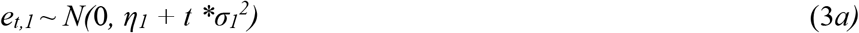

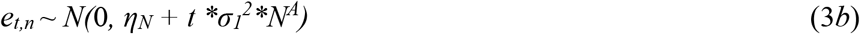

The linear relationship between total accumulated noise and time for both a single and multiple memoranda is illustrated in Fig. 2b.

### Average-then-Diffuse (AtD) Simultaneous model

For this model, the representation of the average is stored as a single particle and diffuses the same as a Perceived point (i.e., a location at which there was a visible stimulus) (See Fig. 2b). Thus, the diffusion term for the representation of a computed average of *N* points, *σ_MN_^2^*, is also *σ_1_^2^*. We do not assume that the representation of the average has the same non-time-dependent noise as a single point, because there could be additional noise from inaccurately averaging multiple points or conversely a reduction in overall noise resulting from the averaging of multiple random variables (constituent points). We denote the variance of the non-time-dependent noise for the Computed mean by *η_MN_*. The difference between the true mean of *N* stimuli and the mean reported at time *t* is therefore,

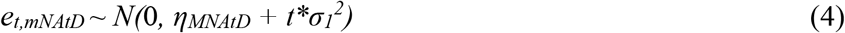

### Diffuse-then-Average (DtA) Simultaneous model

For this model, the individual perceived points are stored as individual, independently diffusing particles and then averaged at the end of the trial. Thus, the diffusion constant of the Computed value is the variance of the average of *N* random variables each with the diffusion constant *σ_1_^2^*\**N^A^*, resulting in an effective diffusion constant for the Computed value of *σ_MN_^2^*=*σ_1_^2^*\**N^A^/N*, where the division by *N* arises from averaging. Again, we allow for a free non-time-dependent-noise term because of the uncertain effects of the averaging calculation itself. For this model, the error in the reported location at time *t* of the average of the mean, *M*, of *N* points, *e_t,MN_*, is:

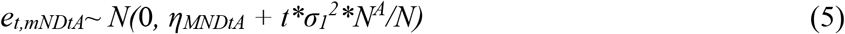

If *A*=1, the AtD and DtA models are identical. We thus used best-fitting values of *A* to help assess model distinguishability for each participant and task condition (see Fig. S7). Also, if *A<1*, then the DtA strategy results in a lower diffusion constant for a Computed value than predicted by the AtD model and results in a smaller average reporting error (see Fig. 2c). If *A*>1, then AtD results in the lower diffusion constant and thus a lower average reporting error. However, we did not find that participants necessarily used the objectively optimal (i.e., lowest-error) strategy in making their decisions.

### Sequentially presented values in working memory

In the Sequential blocks, *N*–1 points were presented immediately (Early points), and the *N*^th^ point was presented halfway through the delay (Late point). Therefore, both our modeling assumes that the diffusion constant for the representation of the *N*–1 early stimuli change with the addition of the *N*^th^ point. In contrast, the representation of the Late stimulus diffuses only for half of the delay time, *T*, as shown in Fig. 2c. We formalize this process by the following model for the report error of the Early (*e_T,NE_*), and Late (*e_T,NL_*) stimuli:

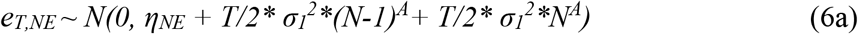

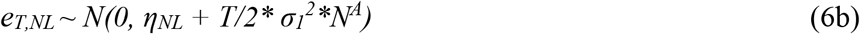

Here *T* is the total time of the delay, and we assumed different non-time-dependent noise for both Early and Late points, *η_NE_* and *η_NL_*, respectively.

### AtD Sequential model

This model assumes that the Early points are averaged immediately and stored as a single point. At *t*=*T*/2, the Late point is presented and the stimulus is immediately combined, through appropriate weighted averaging, with the mean of the Early points. This new mean again diffuses with the same accumulating noise as a single point, as depicted in Fig. 2d. Therefore:

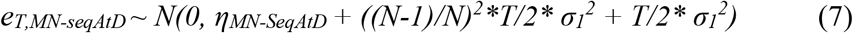

At *t*=*T*/2, the representation of the *N*^th^ stimulus has not accumulated any diffusion noise and only has non-time-dependent noise, which is absorbed in the *η_MN-Seq_* term. The first time-dependent term, *((N-1)/N)^2^*T/2* σ_1_^2^*, results from the appropriate weighted averaging of the mean of the Early points (time-dependent noise of *T/2* σ_1_^2^*) with the Late point (time-dependent noise=0). The final term, *T/2* σ_1_^2^*, is the diffusion of the resultant mean until the end of the delay.

### DtA Sequential model

This model assumes that the representations of all *N* points diffuse as they are presented, resulting in *N*–1 points described by eq. 6a and one point described by eq. 6b. These points are then averaged at the end of the delay, resulting in an overall error of:

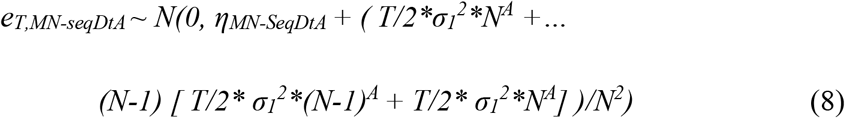

where the constant noise terms from the Early and Late points are absorbed in the *η_MN-SeqDtA_* term, the next term *T/2*σ_1_^2^*N·* is the diffusion in the representation of the last disk shown, and the remaining terms arise from the first *N-1* disks shown. The effect of this averaging on the effective diffusion constant are shown in Fig. 2d.

### Model fitting

To fit both the AtD and DtA models to data from the Perceived and Computed blocks we had to estimate 5 parameters: 1) the non-time-dependent noise of a single point (*η_1_*), 2) the diffusion noise of a single point (*σ_1_^2^*), 3) the non-time-dependent noise of *N* points (*η_N_*), 4) the exponent of storing *N* points (*A*), and 5) the non-time-dependent noise of the mean (*η_MN(AtD or DtA)_*). We fit these models for *N*=2 and *N*=5 separately using trials from the following conditions: Perceived Simultaneous delays of 1, 3, and 6 s, with array sizes 1 (eq. 3a) and *N* (eq. 3b); Computed Simultaneous, delays of 1, 3, and 6 s, with array size *N* (eq. 4 for AtD, eq. 5 for DtA).

Data from the Sequential Perceived and Sequential Computed blocks were fit to the AtD and DtA models with 6 parameters. The additional parameter accounted for differences in the non-time-dependent noise for Early and Late points. We fit these models for *N*=2 and *N*=5 separately using trials from the following conditions: Perceived, delay 1, 3, and 6 s with array sizes 1 (eq. 3a); Sequential Perceived, delay of 3 and 6 s, array size *N* for both Early (eq. 6a) and Late (eq. 6b) points; Sequential Computed, delay of 3 and 6 s, array size *N* (eq. 7 for AtD or eq. 8 for DtA).

Because the mean error for each individual participant was not always zero, when fitting the AtD and DtA models we used the empirical mean error from the condition being fitted as a fixed bias term in the model. Mean error and confidence intervals for each participant for each condition are shown in Fig. S3–6.

We obtained separate maximum-likelihood fits for AtD and DtA models for each individual participant, using the function fmincon in MATLAB to minimize the summed negative log likelihood of obtaining the observed errors for a given condition according to the above equations. Initial parameter values were randomized and the fitting repeated to avoid local minima. Because all models within a given condition had the same number of parameters, we compared log likelihoods to determine the best-fitting model for a given participant. Because the number of parameters are the same, comparing likelihoods produces equivalent model selection to BIC or AIC.

### Goodness-of-fit

To assess how well each participant’s data matched the assumptions of the AtD and DtA models, we also fit a line to the variances of response errors across delays for a given condition for a given participant to obtain empirical estimates of the various diffusion constants (e.g., slope of lines in Fig. 2b; empirical estimate of a Perceived value, 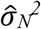, for set size *N*; empirical estimate of a Computed value, 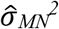, set size *N*). These empirical estimates of the diffusion constants did not enforce the relationships imposed by the AtD and DtA models between the different diffusion constants in each model respectively. We compared the relationships of these empirical estimates of diffusion constants to the relationships assumed by our models.

### AtD Simultaneous expected diffusion constant relationship

Under the AtD hypothesis, for Simultaneous conditions, the Computed mean diffuses with the same diffusion constant as a single value. Thus:

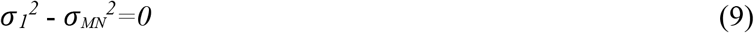

### DtA Simultaneous expected diffusion constant relationship

Under the DtA hypothesis, for Simultaneous conditions, the Computed mean is the average of *N* points each diffusing with a constant of *σ_N_^2^*. Thus:

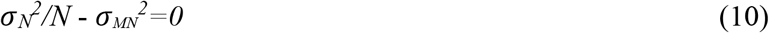

### AtD Sequential expected diffusion constant relationship

Under the AtD hypothesis, for Sequential conditions, the time-dependent noise has variance that increases as *((N-1)/N)^2^*T/2* σ_1_^2^ + T/2* σ_1_^2^)* (eq. 7). Factoring out T gives the diffusion constant for the Computed mean, *σ_MN_^2^*= [(N-1)^2^+N^2^]/(2N^2^*)* σ_1_^2^*. Thus:

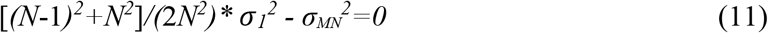

### DtA Sequential expected diffusion constant relationship

Under the DtA hypothesis, for Sequential conditions, the time-dependent noise has variance that increases as *T/2*σ_1_^2^*N^A^ + (N-1) [ T/2* σ_1_^2^*(N-1)^A^ + T/2* σ_1_^2^*N^A^])/N^2^* (eq. 8). By eq. 6a, the diffusion constant for an Early Perceived point, *σ_NE_^2^*, is [0.5* *σ_1_^2^*(N*-1*)^A^ +* 0.5* *σ_1_^2^*N^A^]* and by eq. 6b, the diffusion constant for a Late Perceived point, *σ_NL_^2^*, is *σ_1_^2^*N^A^*. Factoring out T and substituting gives the diffusion constant for the Computed mean, *σ_MN_^2^*= (0.5*σ_NL_^2^+(N*-1*)* σ_NE_^2^)/N^2^*. Thus:

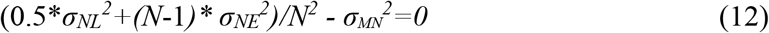

To assess how well the relationships between participant empirical estimates of the diffusion constants matched these assumptions, for each participant we simulated 1000 iterations of a participant performing the task using the best-fit model for the given true participant and the maximum likelihood estimate parameters for that participant. We then estimated the empirical diffusion constants for each of these iterations as we did for our participants, namely fitting a line to the measured variance of the simulated errors across delays, for each condition and iteration. Our 1000 simulations gave us an expected range around the expected diffusion constant relationships detailed in eq. 9–12 to compare to our participants’ empirical diffusion constant relationships. Participants whose empirical diffusion constant relationships fell within the central 95% of the simulated expected range were considered well fit by their model.

## Acknowledgements

We thank Adrian Radillo and Gaia Tavoni for their discussions early in the development of this project, particularly regarding early formulations of the models used here and task structure.

## Supplemental Figures

**Supplemental Figure S1:**
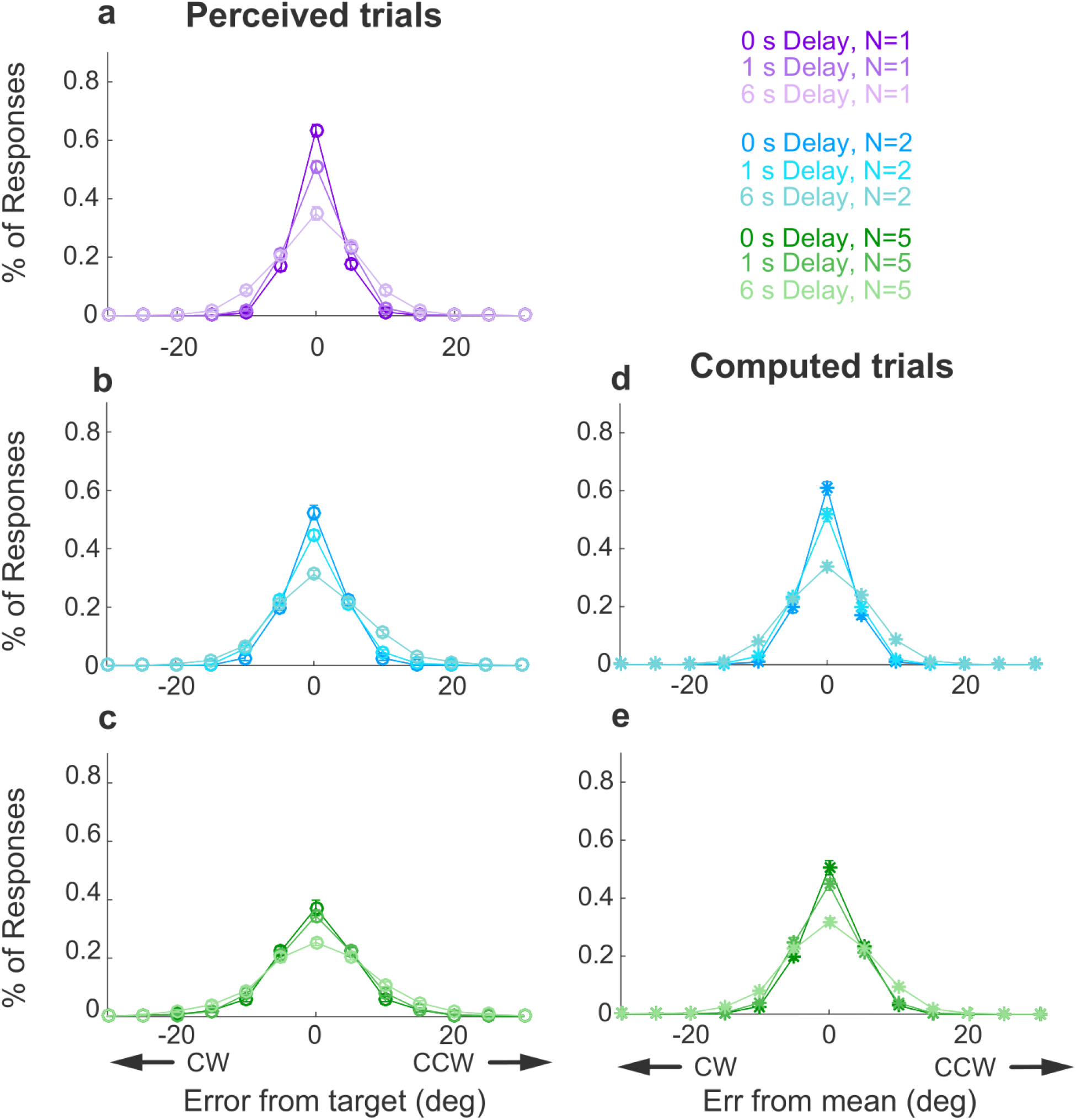
Full error distributions in Simultaneous conditions. Each panel shows a histogram of mean error for different delays (colors, as indicated) for Perceived trials (left column: **a**) set size=1; ***b***) set size=2; ***c***) set size=5) and for Computed trials (right column; ***d***) set size=2; ***e***) set size=5). In each panel, points and error bars are mean±SEM across all participants. Note that in all cases, 95% of the distributions fall between −30° and 30°, justifying our exclusion of larger errors as off-target responses.

**Supplemental Figure S2:**
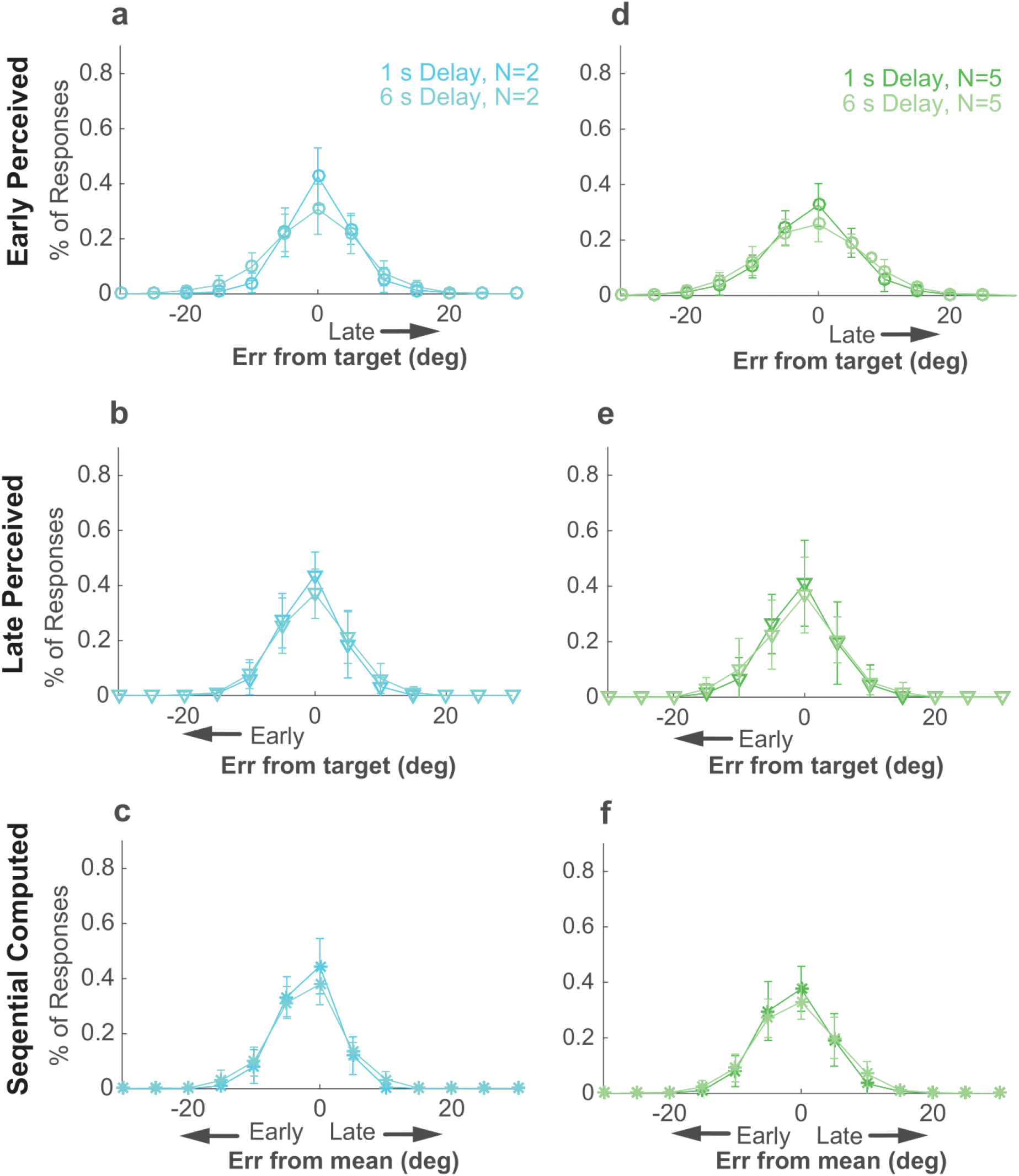
Full error distributions in Sequential conditions. **a**) Histogram of mean Early Perceptual error for different delays for set size 2. **b**) Histogram of mean Late Perceptual error for different delays for set size 2. **c**) Histogram of mean Computed error for different delays for set size 2. **d-f**) As in **a-c** but for set size 5. In each panel, points and error bars are mean±SEM across participants. Note that in all cases, 95% of the distributions fall between −30° and 30°, justifying our exclusion of larger errors as off-target responses.

**Supplemental Figure S3:**
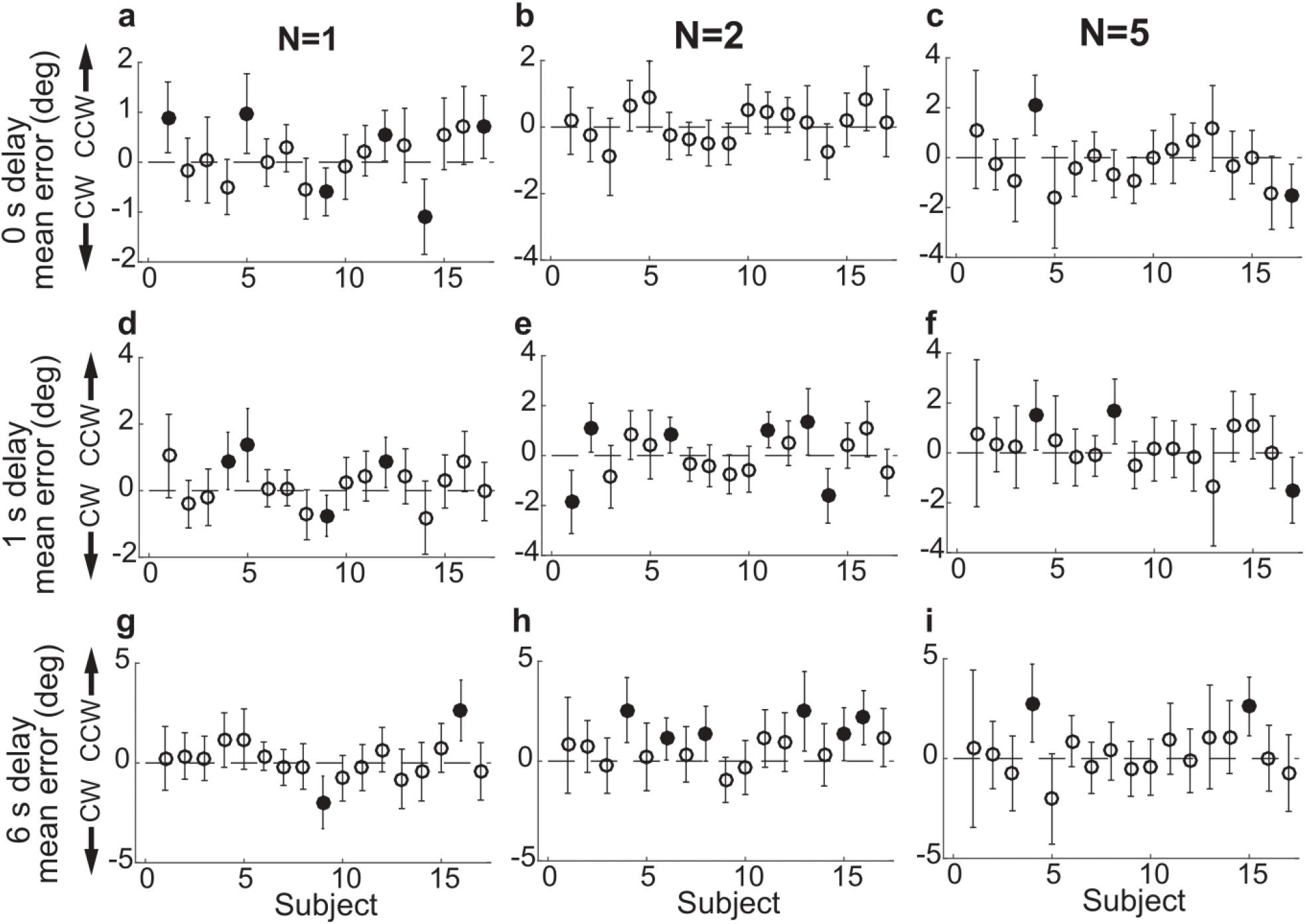
Subject-wise mean response error in the Simultaneous Perceived condition. **a**) Delay=0 s, set size=1. **b)** Delay=0, set size=2. **c)** Delay=0 s, set size =5. **d–f)** as in **a–c** but for delay of 1 s. **g–i)** as in **a–c** but for delay of 6 s. In all panels, errorbars are ±95% confidence intervals. Filled points indicate that 0 is not included in the confidence interval; i.e., there is bias in subject errors.

**Supplemental Figure S4:**
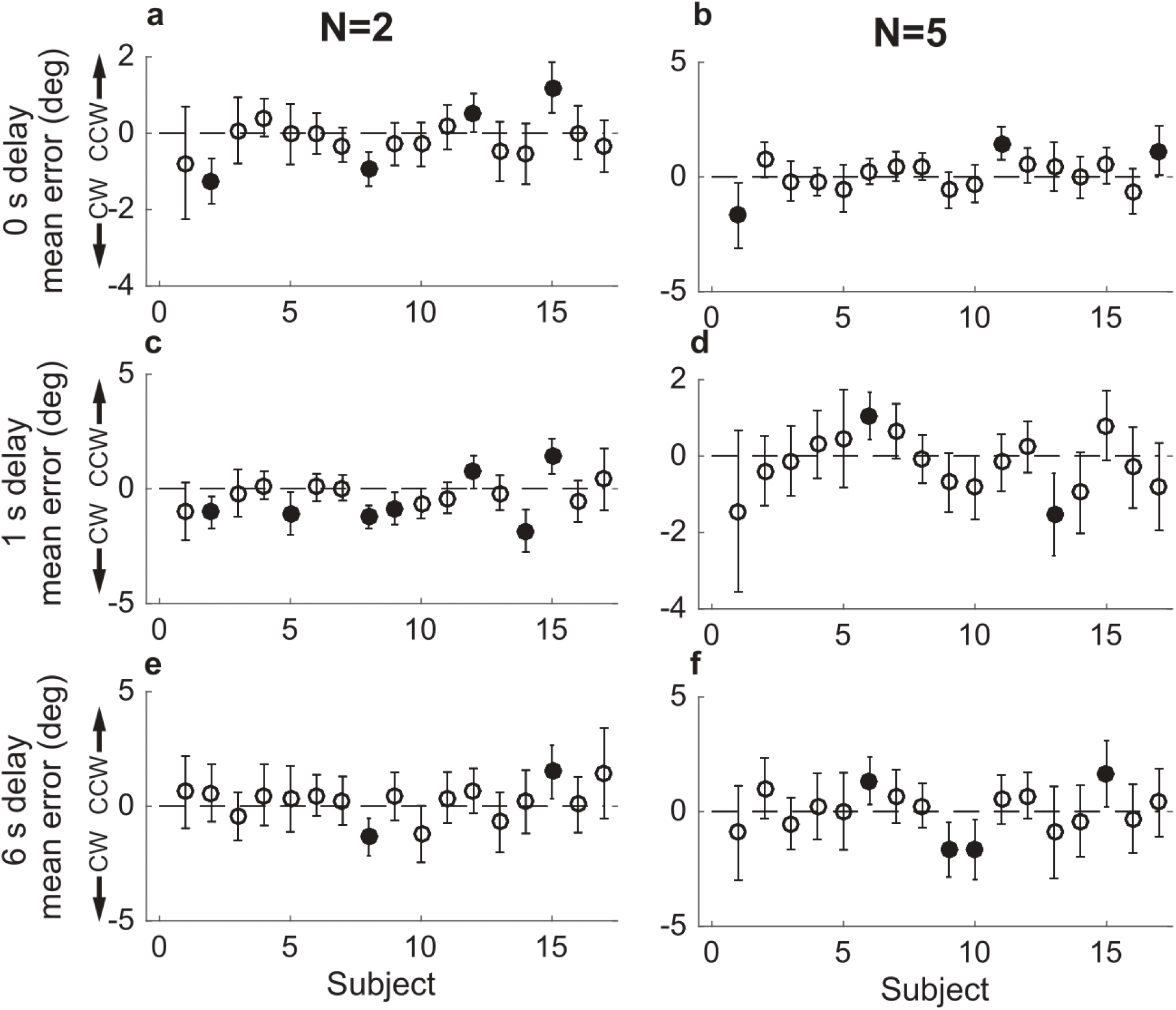
Subject-wise mean error in the Simultaneous Computed condition. **a**) Delay=0 s, set size=2. **b)** Delay of 0, set size=5. **c–d)** as in **a–b** but for delay of 1 s. **e-–f)** as in **a–b** but for delay of 6 s. In all panels, errorbars are ±95% confidence intervals. Filled points indicate that 0 is not included in the confidence interval; i.e., there is bias in subject errors.

**Supplemental Figure S5:**
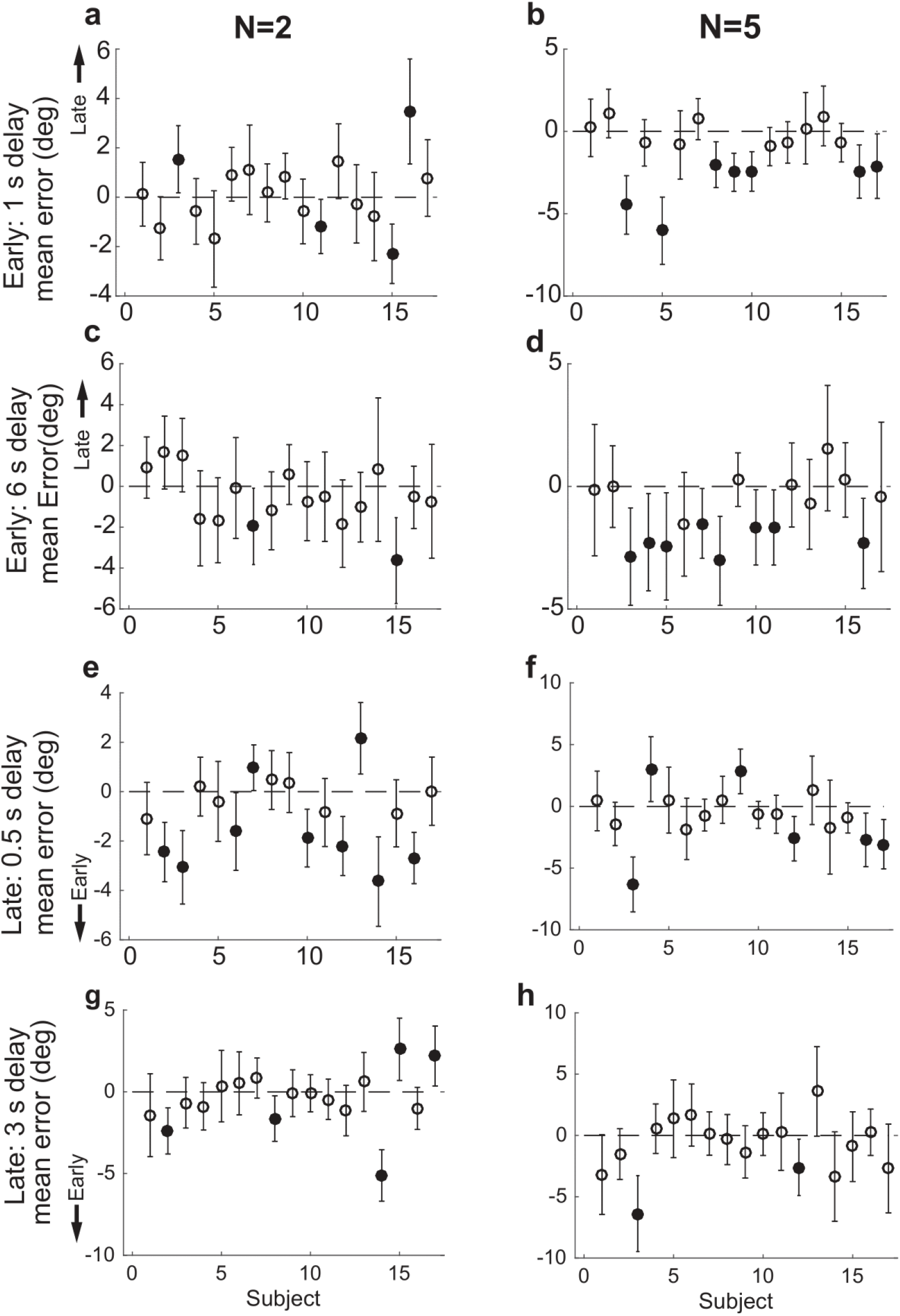
Subject-wise mean error in the Sequential Perceived condition. **a**) Delay=1 s, set size=2 for Early samples. **b)** Delay=1, set size=5 for Early samples. **c–d)** as in **a–b** but for delay of 6 s. **e**) Delay=0.5 s, set size=2 for Late samples. **f)** Delay=0.5 s, set size=5 for Late samples. **g–h)** as in **e–f** but for delay of 3 s. In all panels, errorbars are ±95% confidence intervals. Filled points indicate that 0 is not included in the confidence interval; i.e., there is bias in subject errors.

**Supplemental Figure S6:**
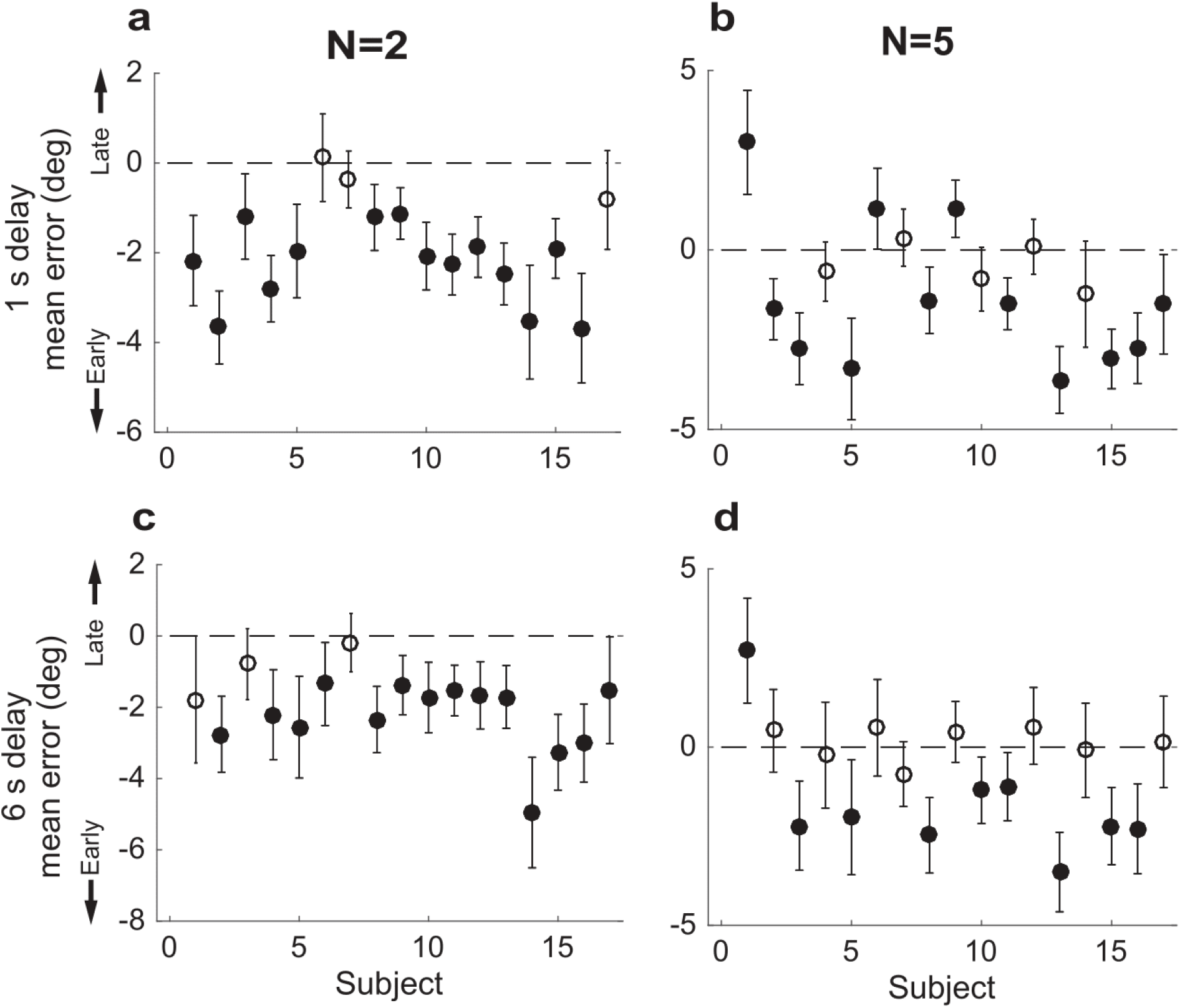
Subject-wise mean error in the Sequential Computed condition. **a**) Delay=1 s, set size=2. **b)** Delay=1, set size=5. **c–d)** as in **a–b** but for delay of 6 s. In all panels, errorbars are ±95% confidence intervals. Filled points indicate that 0 is not included in the confidence interval; i.e., there is bias in subject errors (which in this case tended to be towards the mean computed from the early points).

**Supplemental Figure S7:**
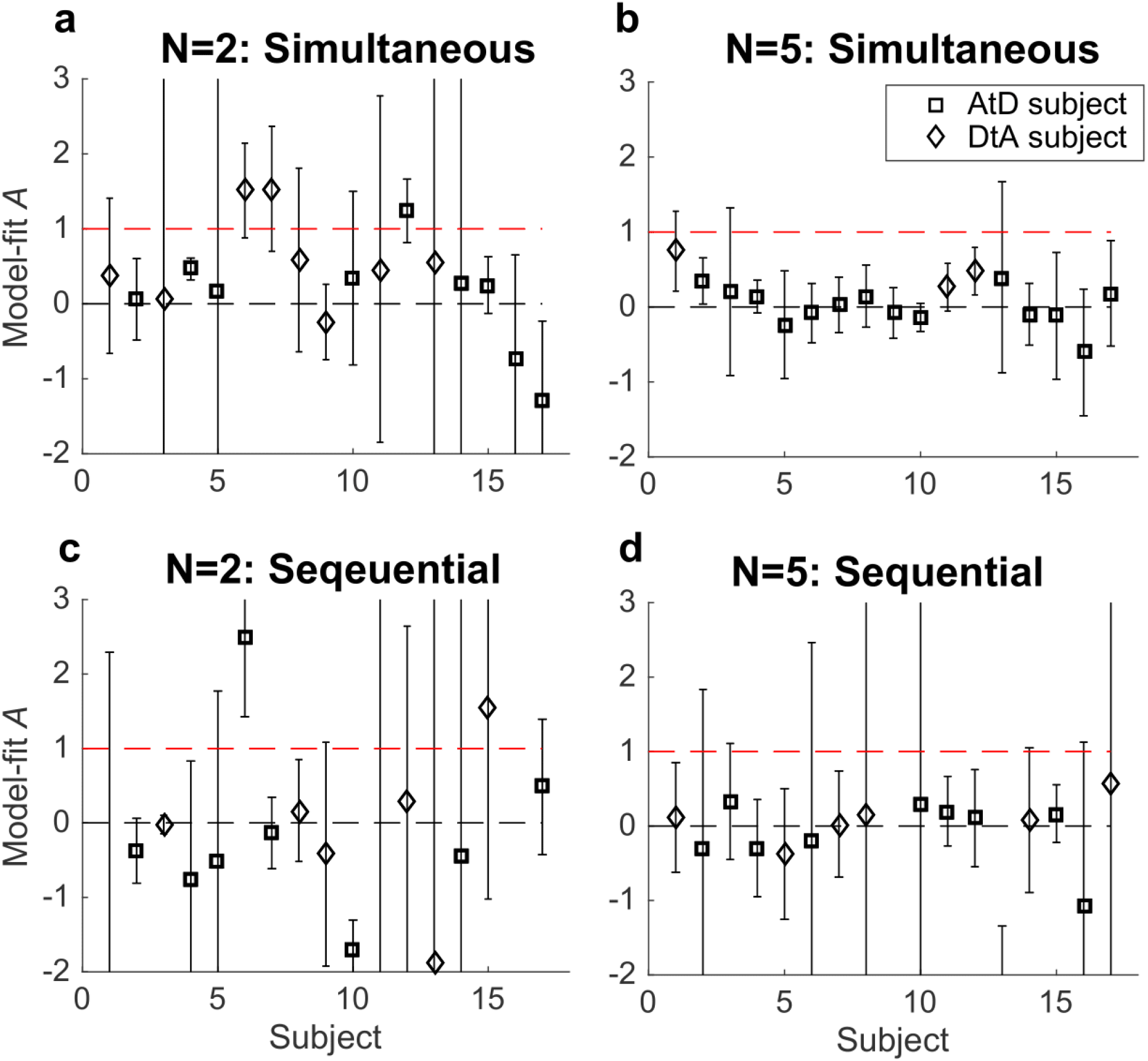
Subject-based estimates of *A*. **a,b**) Simultaneous condition. **a**) Model-fits for *A* for AtD (square) and DtA (diamond) participants at set size 2. **b**). Model-fits for *A* for all participants at set size 5. **c,d**) Same as **a,b** but for the Sequential condition. In all panels, errorbars are ±95% confidence intervals based on the Hessian computed during model fitting. Note that *A*=0 implies no difference between the diffusion constant for a single and *N* points, whereas *A*=1 implies that the variance and diffusion constant relationship predictions of the AtD and DtA models are equal and thus the models cannot be distinguished from each other.

**Supplemental Figure S8:**
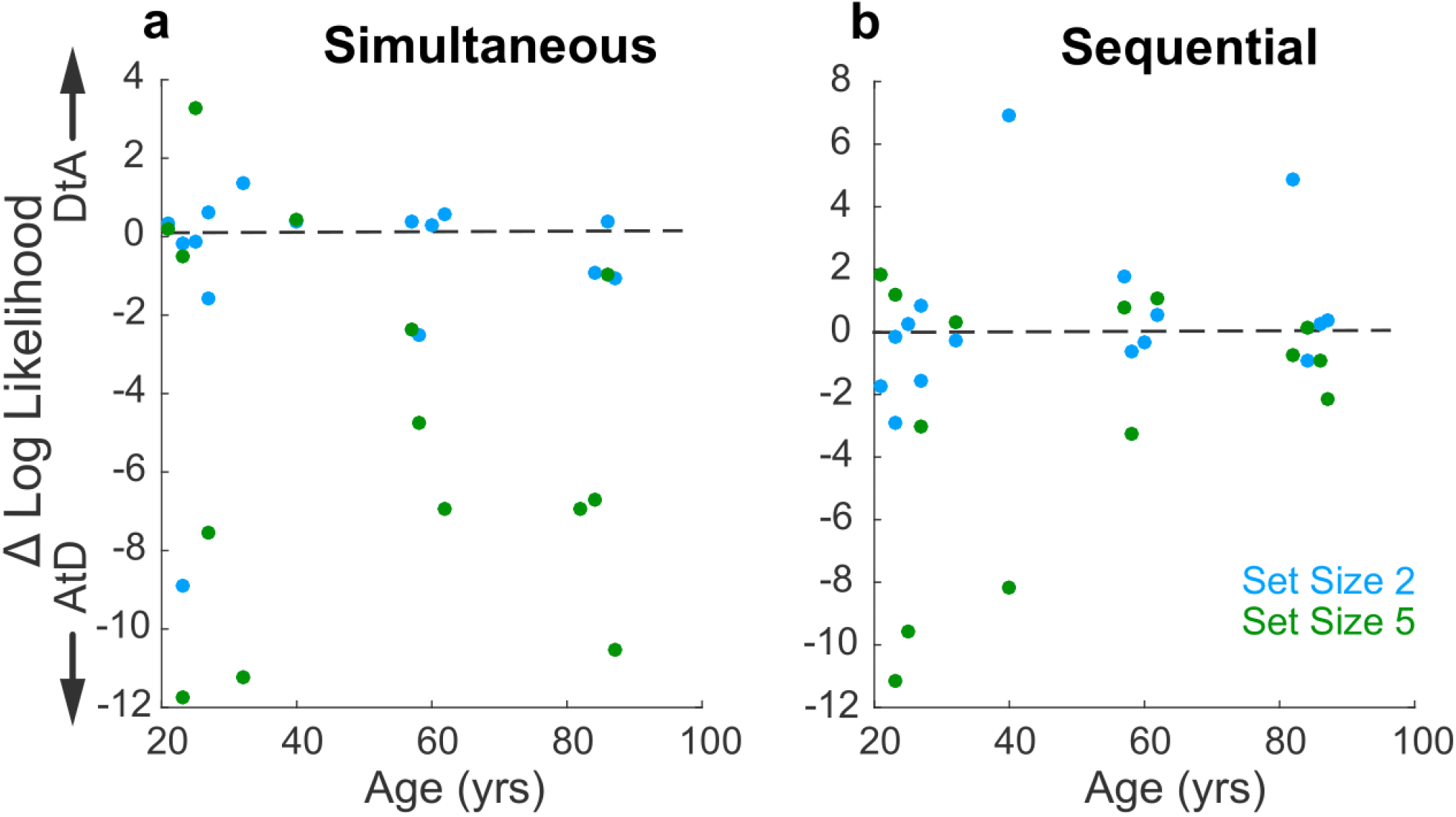
Relationship between log likelihood difference for the two strategies and age. **a)** Log likelihood comparison for AtD and DtA (negative favors AtD) for set sizes 2 and 5 under Simultaneous conditions is not dependent upon age (correlation, *ps*>0.20 computed separately for each set size). **b)** Same as in A, but for the Sequential Conditions (*ps*>0.20).

